# Extended genomic analyses of the broad-host-range phages vB_KmiM-2Di and vB_KmiM-4Dii reveal slopekviruses have highly conserved genomes

**DOI:** 10.1101/2022.04.06.486684

**Authors:** Thomas Smith-Zaitlik, Preetha Shibu, Anne L. McCartney, Geoffrey Foster, Lesley Hoyles, David Negus

**Affiliations:** Department of Biosciences, Nottingham Trent University, UK; Life Sciences, University of Westminster, UK; Department of Food and Nutritional Sciences, University of Reading, UK; SRUC Veterinary Services, Inverness, UK

**Keywords:** *Slopekvirus*, phage diversity, lytic phage, myovirus, pangenome, *Klebsiella oxytoca* complex, homing endonuclease

## Abstract

High levels of antimicrobial resistance among members of the *Klebsiella oxytoca* complex (KoC) have led to renewed interest in the use of bacteriophage (phage) therapy to tackle infections caused by these bacteria. In this study we characterized two lytic phages, vB_KmiM-2Di and vB_KmiM-4Dii, that were isolated from sewage water against two GES-5-positive *Klebsiella michiganensis* strains (PS_Koxy2 and PS_Koxy4, respectively). ViPTree analysis showed both phages belonged to the genus *Slopekvirus. rpoB* gene-based sequence analysis of 108 presumptive *K. oxytoca* isolates (*n*=59 clinical, *n*=49 veterinary) found *K. michiganensis* to be more prevalent (46 % clinical and 43 % veterinary, respectively) than *K. oxytoca* (40 % clinical and 6 % veterinary, respectively). Host range analysis against these 108 isolates found both vB_KmiM-2Di and vB_KmiM-4Dii showed broad lytic activity against KoC species. Several putative homing endonuclease genes were encoded within the genomes of both phages, which may contribute to their broad host range. Pangenome analysis of 24 slopekviruses found that genomes within this genus are highly conserved, with more than 50 % of all predicted coding sequences representing core genes at ≥95 % identity and ≥70 % coverage. Given their broad host ranges, our results suggest vB_KmiM-2Di and vB_KmiM-4Dii represent attractive potential therapeutics. In addition, current recommendations for phage-based pangenome analyses may require revision.

## INTRODUCTION

Members of the *Klebsiella oxytoca* complex (KoC) are divided into phylogroups based on the sequence of their chromosomally encoded β-lactamase (*bla*_OXY_) gene. The current phylogroups are *Klebsiella michiganensis* (KoI, with KoV sub-lineage), *K. oxytoca* (KoII), *K. spallanzanii* (KoIII), *K. pasteurii* (KoIV), *K. grimontii* (KoVI) and *K. huaxiensis* (KoVIII). KoVII has been described based on a single isolate (1–3).

Several members of the KoC can cause a variety of infections in humans including urinary tract infections (UTIs), septicaemia and *Clostridioides*-negative antibiotic-associated haemorrhagic colitis (AAHC) (4–6). The rapid development of antimicrobial resistance (AMR) and the lack of novel antibiotics is a serious public health concern. Of 41 strains of the KoC isolated from bloodstream infections in the UK and Ireland, 100 % were phenotypically resistant to amoxicillin and cefuroxime, 75.6 % to piperacillin-tazobactam, 73.2 % to amoxicillin-clavulanate and 48.8 % to ciprofloxacin (7). In a survey of 5,724 clinical isolates of *K. oxytoca*, the SENTRY Antimicrobial Surveillance Program identified rates of non-susceptibility of *K. oxytoca* to various antibiotics: 1.8 % carbapenems, 12.5 % ceftriaxone, 7.1 % ciprofloxacin, 0.8 % colistin and 0.1 % tigecycline (8). GES-positive clinical strains of the KoC have also been identified recently (9,10).

KoC bacteria can also cause disease in animals. Of 336 samples collected from companion animals, 11 (3.3 %) isolates were identified as *K. oxytoca*. These were typically recovered from the urogenital system and 81.8 % were resistant to ampicillin (11). *Klebsiella* spp. were detected in 51/1541 (3.3 %) equine samples. Two *K. michiganensis* and one *K. oxytoca* were identified as non-repetitive cefotaxime-resistant isolates. These three isolates were phenotypically resistant to gentamicin, tobramycin, tetracycline, doxycycline, chloramphenicol and trimethoprim/sulfamethoxazole (12).

Accurate identification at the species level is important for recognising the epidemiological and clinical significance of each member of the KoC in both humans and animals. Current diagnostics are unable to consistently differentiate between members of the KoC leading to historic misidentification of non-*K. oxytoca* strains as *K. oxytoca* (10). The phenotypic similarity of the complex members prevents accurate identification using biochemical tests such as API 20E tests (1). While matrix-assisted laser desorption/ionisation-time of flight mass spectrometry (MALDI-TOF MS) is a rapid and cost-effective diagnostic tool, many databases in clinical and veterinary use have not been updated and lack biomarkers required to consistently differentiate members of the KoC (13,14). The use of 16S rRNA gene sequencing is considered unsuitable for identifying *Klebsiella* at the species level as the gene sequence is so highly conserved in this taxon (15). *gyrA* and *rpoB* gene sequences have been used to distinguish *K. grimontii* and *K. huaxiensis* from other members of the KoC, supported by results from average nucleotide identity (ANI) analysis of genome sequence data (16,17). More recently, *rpoB* gene sequence analysis has been used to identify KoC, *Klebsiella pneumoniae* complex and *Raoultella* spp. isolates recovered from the faeces of healthy women and breast-fed infants (18). Multi-locus sequence typing (MLST) can also be used to assign strains to species of the KoC (10).

Bacteriophages (phages) are promising alternatives or adjuncts to current antibiotic therapies. We previously published an extensive review on *Klebsiella* phage and their potential as therapeutics (19). The majority of *Klebsiella* phage publications focus on *K. pneumoniae*. Phage vB_Kox_ZX8 was isolated from human faeces; this phage was shown to clear bacteraemia caused by a clinical strain of *K. oxytoca* in BALB/c mice (20). Another recent study described 30 novel phages that were active against *Klebsiella* species including KoC members *K. oxytoca* and *K. michiganensis* (21). Of the phages isolated, 15 were active against the single *K. michiganensis* strain tested, whereas 16 showed activity against at least one of the five *K. oxytoca* strains tested. More recently, the lytic *Drexlerviridae* phage KMI8 was isolated against *K. michiganensis* (22). KMI8 was lytic against 3/5 *K. michiganensis* strains but not *K. pneumoniae* (0/5) or *K. oxytoca* (0/1). To the best of our knowledge, ISF3 and ISF6 (23) and RP180 (24) are the only three *Raoultella* phages recorded in the literature. They were isolated against *Raoultella ornithinolytica* and *Raoultella* spp., respectively.

This study aimed to characterise the morphology, genomes and host ranges of two lytic phages, vB_KmiM-2Di and vB_KmiM-4Dii, isolated against two strains of GES-5-encoding *K. michiganensis* (PS_Koxy2 and PS_Koxy4) (25). We used *rpoB* gene sequence analysis to accurately identify 108 clinical and veterinary isolates identified as *K. oxytoca* using MALDI-TOF MS and/or API 20E. These isolates were used in our host range analysis, to determine the therapeutic potential of vB_KmiM-2Di and vB_KmiM-4Dii against *Klebsiella* spp. A pangenome analysis was undertaken to compare our new phage genomes with those of their closest relatives.

## METHODS

### Strain information

Details of all strains included in this study can be found in **Table 1.**

**Table 1.**
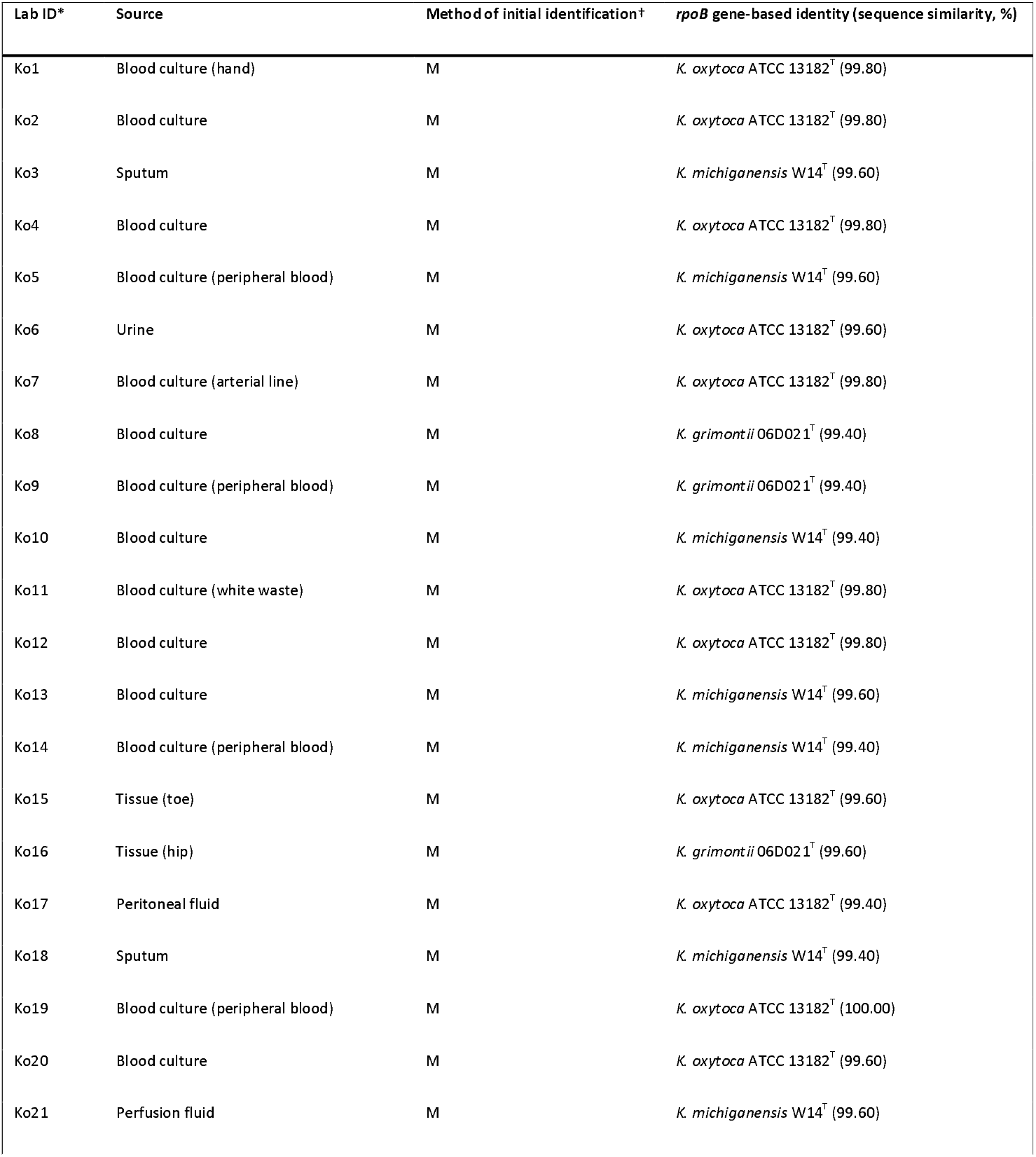
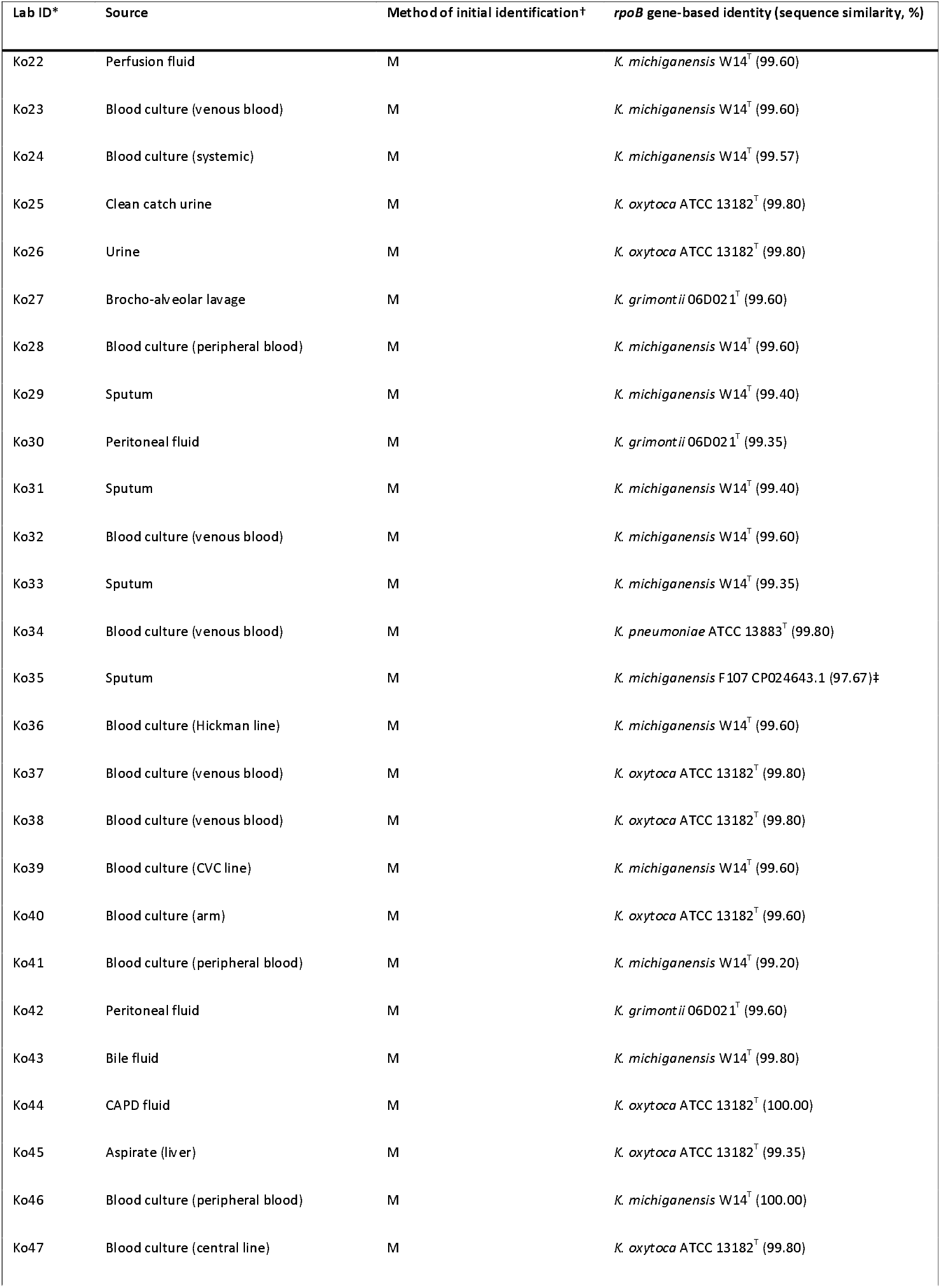
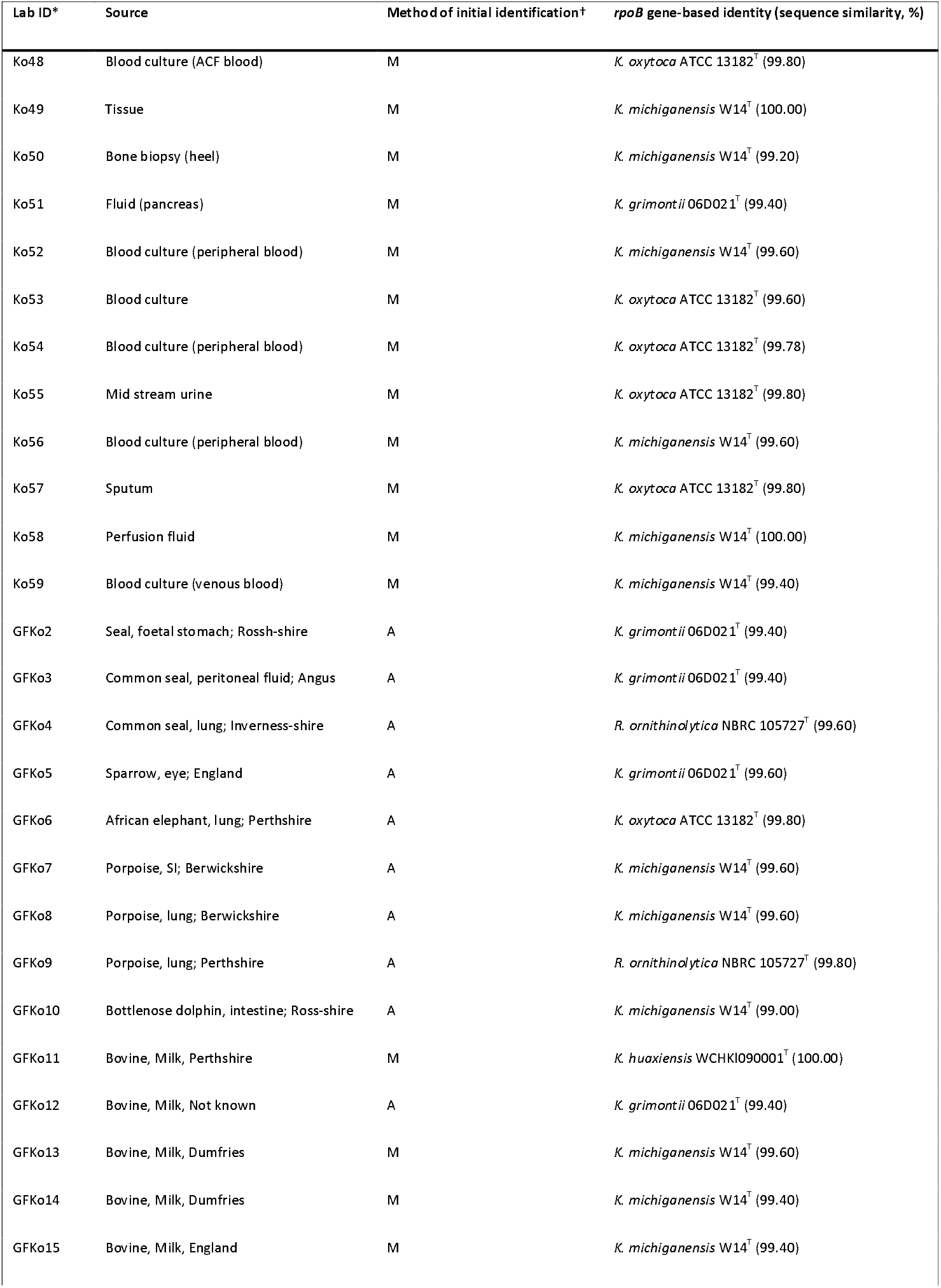
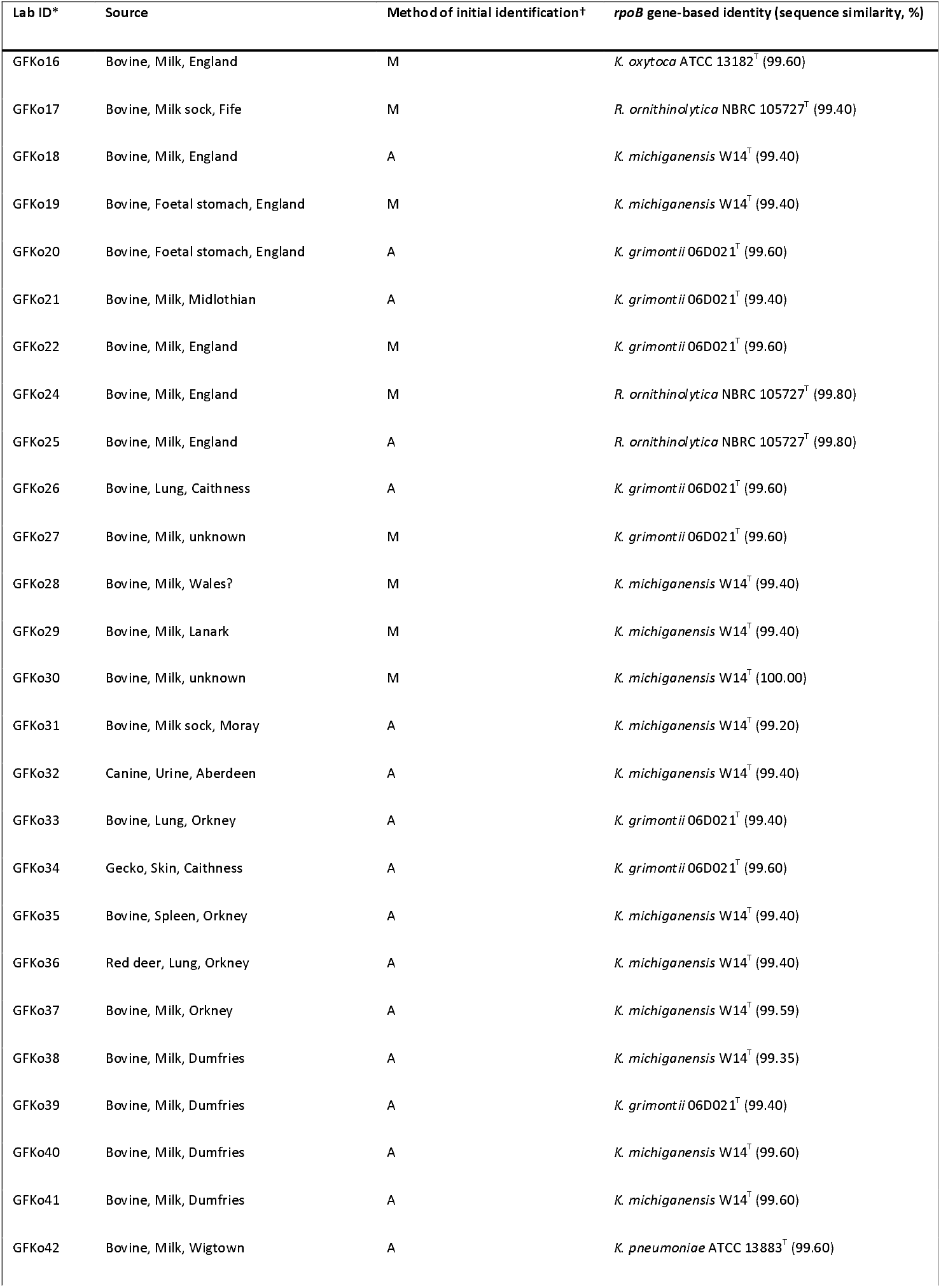
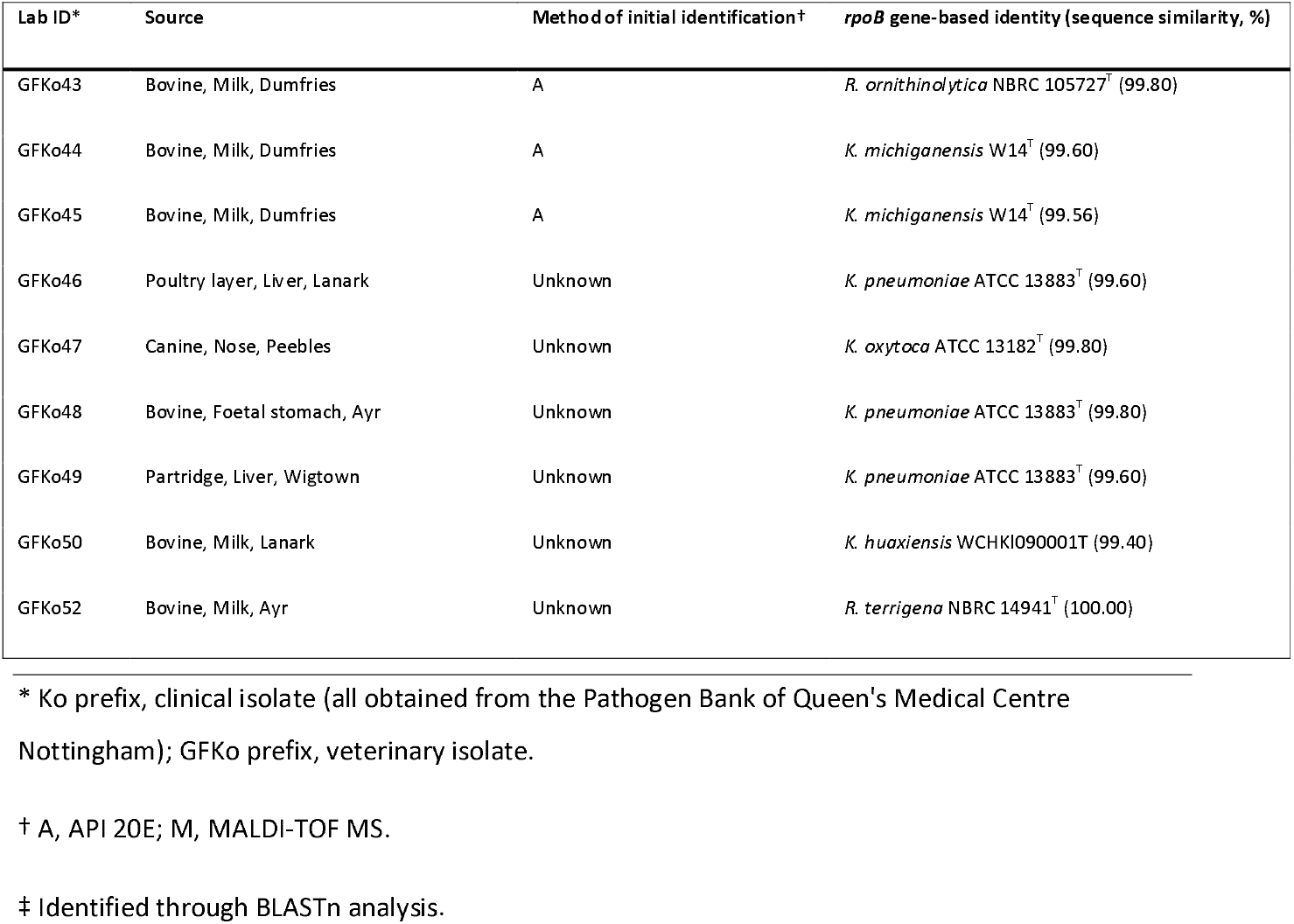
Clinical and veterinary isolates included in this study. With the exceptions of Ko8, Ko31 and GFKo17 (*K. oxytoca/R. ornithinolytica*), all strains were initially identified as *K. oxytoca*.

### Isolation of lytic phage

Filter-sterilised sewage samples (0.45 μm cellulose acetate filter; Millipore) collected from mixed-liquor tanks at Mogden Sewage Treatment Works (March 2017) were screened against *K. michiganensis* strains PS_Koxy2 and PS_Koxy4 (10,25). Firstly, 9 mL of filter-sterilised sewage were added to 1 mL of 10× concentrated sterile nutrient broth (NB) (Oxoid Ltd) containing 50 mM CaCl_2_ and 50 mM MgCl_2_. This was then inoculated with 200 μL of overnight culture from each strain and incubated for 6 h at 37 °C. The samples were centrifuged at 10,000 rpm for 5 min; the supernatants were aliquoted (200 μL) and used in spot assays to identify lytic phage. Plaques were propagated to purity to create phage stocks.

### Growth media and culture conditions

Bacterial cultures were initially streaked onto MacConkey agar (Sigma Aldrich) to ensure purity before being grown on nutrient agar (NA) (Sigma Aldrich). NB (Sigma Aldrich) was used for overnight cultures, incubated aerobically at 37 °C. All media used for phage assays were supplemented with CaCl_2_ and MgCl_2_ (final concentration 0.5 mM) unless otherwise specified.

### Colony PCR and sequencing of *rpoB* gene products

Forward (5’-GTTTTCCCAGTCACGACGTTGTAGGCGAAATGGCGGAAAACCA-3’) and reverse (5’-TTGTGAGCGGATAACAATTTCGAGTCTTCGAAGTTGTAACC-3’) *rpoB*-specific primers (Macrogen) (23) were diluted to 10 μM in DNase-free H_2_O. A single colony for each isolate was touched using a sterile loop and dipped into the PCR master mix: 25 μL MangoMix™ (Meridian Bioscience); forward and reverse primers (0.5 μM final concentration); 20 μL DNase-free H_2_O. DNA loading dye was included in the MangoMix™. The positive control tube contained 2 μL of concentrated *K. michiganensis* PS_Koxy1 DNA (10) and the negative control contained master mix alone (no template DNA). The cycle conditions were: initial denaturation, 95 °C for 10 min; 35 cycles 95 °C for 30 s, 54 °C for 30 s, 72 °C for 1 min; final extension, 72 °C for 5 min. PCR products were checked for single bands of expected size (1076 nt) using agarose gel electrophoresis (1 % agarose gel in 1x TAE buffer; 100 V, 40 min) against a GeneRuler 1 kb ladder (ThermoFisher Scientific).

The Thermo Scientific™ Gene JET PCR Purification Kit was used to clean PCR products. Purified samples were checked for DNA concentration and purity using a NanoDrop™ 2000 spectrophotometer (24). Samples were then adjusted to 10 ng/μL and sent for sequencing (Source BioScience) using the MLST forward primer (5’-GTTTTCCCAGTCACGACGTTGTA-3’) (25).

### Phylogenetic analysis of *rpoB* gene sequences

Returned *rpoB* gene sequences were trimmed to 501 nt using Geneious Prime (v2020.0.5) by extracting the sequence between 276 and 776 nt, and with reference to the 45 *rpoB* allele sequences (released 19 June 2020) available for download from the PubMLST *Klebsiella oxytoca/michiganensis/grimontii* typing database (26). *rpoB* gene sequences were extracted from the following genomes and used in analyses: *K. oxytoca* (GCA_900977765), *K. spallanzanii* (GCA_901563875), *K. pasteurii* (GCA_901563825), *K. grimontii* (GCA_900200035), *K. michiganensis* (GCA_901556995), *K. pneumoniae* subsp. *ozaenae* (GCA_000826585), *K. pneumoniae* subsp. *rhinoscleromatis* (GCA_000163455), *K. pneumoniae* subsp. *pneumoniae* (GCA_000742135), *K. quasipneumoniae* subsp. *similipneumoniae* (GCA_900978135), *K. quasipneumoniae* subsp. *quasipneumoniae* (GCA_000751755), *K. africana* (GCA_900978845), *‘K. quasivariicola’* (GCA_000523395), *K. variicola* subsp. *tropica* (GCA_900978675), *K. variicola* subsp. *variicola* (GCA_900977835), *K. aerogenes* (GCA_003417445), *K. indica* (GCA_005860775) and *K. huaxiensis* (GCA_003261575). *Raoultella electrica* (GCA_006711645), *R. terrigena* (GCA_006539725), *R. planticola* (GCA_000735435) and *R. ornithinolytica* (GCA_001598295) *rpoB* gene sequences were also included in analyses, as these taxa should be classified as *Klebsiella* spp. (27). A multiple-sequence alignment (MSA; available as **Supplementary Material**) was created using Clustal Omega (v1.2.2). The Jukes–Cantor genetic distance model was used to generate a neighbour-joining tree using the *rpoB* gene sequence of *K. aerogenes* ATCC 13048^⊤^ as an outgroup. The resulting newick file (available as **Supplementary Material)** was exported to iTOL (https://itol.embl.de/) (v6.1.1) (28)) for visualisation and annotation of the phylogenetic tree.

### Host range analysis

Sterile molten top NA (3 mL; 0.2 % SeaPlaque Agarose, Lonza) supplemented with CaCl_2_ and MgCl_2_ (both at 5 mM) was aliquoted into sterile test tubes held at 45 °C. Each tube was then inoculated with 250 μL of an overnight culture of the prospective host strain, and gently swirled to mix the contents before being poured onto an NA plate. The plate was gently swirled to ensure even distribution of top agar. Once set, 5 μL aliquots of both phages were spotted onto the plate. Plates were incubated overnight at 37 °C. Next day, plates were inspected for lysis, with results recorded according to a modification of Haines *et al.* (32): ++, complete lysis; +, hazy lysis; 0, no visible plaques. We also noted whether depolymerase activity (d) was evident (i.e., formation of haloes around plaques).

### Phage concentration

The Vivaspin 20 50 kDa centrifugal concentrator (Cytiva) was used to concentrate 20 mL of filter-sterilised propagated phage. Samples were spun at 3000 ***g*** until only 200 μL of sample remained. This concentrated phage stock was stored at 4 °C.

### TEM

Formvar/carbon-coated 200 mesh copper grids (Agar Scientific) were prepared via glow discharge (10 mA, 10 s) using a Q150R ES sputter coater (Quorum Technologies Ltd). Phage suspensions (15 μL) were pipetted onto the grid surface for 30 s before removal using filter paper. Samples were stained using 15 μL of 2 % phosphotungstic acid. Excess stain was removed using filter paper and grids were air-dried. Samples were visualized using a JEOL JEM-2100Plus (JEOL Ltd) TEM and an accelerating voltage of 200 kV. Images were analysed and annotated using ImageJ (https://imagej.net/Fiji).

### Phage DNA extraction

Nuclease-free H_2_O was added to 200 μL of concentrated phage for a total final volume of 450 μL. This was then incubated at 37 °C for 1.5 h with 50 μL of 10× CutSmart Buffer (New England BioLabs) supplemented with 5 mM CaCl_2_, 10 μL of DNase I (1 U/μL) (Thermo Scientific) and RNase A (10 mg/mL) (Thermo Scientific). Next, 20 μL of EDTA (final concentration 20 mM) and 1.3 μL of Proteinase K (20 mg/mL) (Qiagen) were added and incubated at 56 °C for 1.5 h. The Qiagen DNeasy Blood & Tissue Kit (Qiagen) was used to extract and purify the phage DNA. DNA was eluted in 20 μL of the kit’s AE buffer. Phage DNA integrity was checked using agarose gel electrophoresis (1 % agarose gel in 1× TAE buffer; 70 V, 90 min) against a GeneRuler 1 kb ladder (ThermoFisher Scientific).

### Phage DNA sequencing, genome assembly and characterization

Sequence data were generated on our in-house Illumina MiSeq platform. Extracted DNA was adjusted to a concentration of 0.2 ng/μL and treated using the Nextera XT DNA library preparation kit (Illumina) to produce fragments of approximately 500 bp. Fragmented and indexed samples were run on the sequencer using a Micro flow cell with the MiSeq Reagent Kit v2 (Illumina; 150-bp paired-end reads) following Illumina’s recommended denaturation and loading procedures. Quality of raw sequence data was assessed using FastQC v0.11.9. Reads had a mean phred score above 30 and contained no adapter sequences, so data were not trimmed. Genomes were assembled using SPAdes v3.13.0 (33), and visualized using Bandage v0.8.1 (34). Contamination and completeness of genomes were determined using CheckV v0.8.1 (35). Genomes were screened for antimicrobial resistance genes using the Resistance Gene Identifier (v5.2.0) of the Comprehensive Antibiotic Resistance Database (v3.1.4) (36).

ViPTree v1.9 (37) was used to determine whether the phage genomes were closely related to previously described double-stranded DNA viruses. Initial analyses showed them to be closely related to *Klebsiella virus KP15* (genus *Slopekvirus*) (38). Other *Slopekvirus* genomes were identified in GenBank and from the literature **(Table 2)** and included in a second ViPTree analysis. All publicly available genome sequences were also compared against the genomes of *Klebsiella* phage vB_KmiM-2Di and *Klebsiella* phage vB_KmiM-4Dii using pyani v0.2.11 (ANIm) (39).

**Table 2.** *Slopekvirus* genomes included in this study

| Phage | Genome<br>accession | Genome<br>size (nt) | CDS* | CheckV |  |  | Reference |
| --- | --- | --- | --- | --- | --- | --- | --- |
|  |  |  |  | Quality† | Completeness (%) | Contamination (%) |  |
| *Klebsiella* phage vB_KmiM-2Di | MZ707156 | 177200 | 275 | High-quality | 99.34 | 0 | This study |
| *Klebsiella* phage vB_KmiM-4Dii | MZ707157 | 174857 | 272 | High-quality | 98.02 | 0 | This study |
| *Klebsiella virus* KP15 | NC_014036 | 174436 | 269 | High-quality | 97.79 | 0 | (63) |
| *Klebsiella virus* KP27 | NC_020080 | 174413 | 271 | High-quality | 97.78 | 0 | (63) |
| *Klebsiella virus* Matisse | NC_028750 | 176081 | 273 | High-quality | 98.73 | 0 | (66) |
| *Escherichia* phage phT4A | NC_055712 | 171598 | 259 | High-quality | 96.20 | 0 | (67) |
| *Enterobacter virus* Eap3 | NC_041980 | 175814 | 275 | High-quality | 98.56 | 0 | (68) |
| *Klebsiella virus* Miro | NC_041981 | 176055 | 274 | High-quality | 98.70 | 0 | (69) |
| *Klebsiella virus* PMBT1 | NC_042138 | 175206 | 274 | High-quality | 98.22 | 0 | (70) |
| *Klebsiella* phage vB_KpM-KaiD | LR881140 | 174351 | 268 | Complete | 100 | 0 | (21) |
| *Klebsiella* phage vB_KpM-SoFaint | LR881141 | 175933 | 271 | Complete | 100 | 0 | (21) |
| *Klebsiella* phage vB_KpM-Milk | LR881142 | 176734 | 271 | Complete | 100 | 0 | (21) |
| *Klebsiella* phage vB_KoM-Liquor | LR881143 | 176734 | 270 | Complete | 100 | 0 | (21) |
| *Klebsiella* phage vB_KoM-Pickle | LR881145 | 175221 | 273 | High-quality | 98.23 | 0 | (21) |
|  |  |  |  | Quality† | Completeness (%) | Contamination (%) |  |
| Klebsiella phage vB_KpM-Mild | LR881147 | 176856 | 271 | Complete | 100 | 0 | (21) |
| Klebsiella phage vB_KoM-MeTiny | LR883651 | 175419 | 272 | Complete | 100 | 0 | (21) |
| Klebsiella phage vB_KpnM_05F‡ | LR746310 | 174231 | 272 | Complete | 100 | 0 | (32) |
| Klebsiella phage KOX8‡ | MN101221 | 131200 | 190 | Medium-quality | 73.55 | 0 | Unpublished |
| Klebsiella phage KOX10‡ | MN101223 | 168074 | 252 | High-quality | 94.22 | 0 | Unpublished |
| Klebsiella phage KOX11‡ | MN101224 | 59583 | 88 | Low-quality | 33.42 | 0 | Unpublished |
| Klebsiella phage vB_KpnM_P-KP2‡ | MT157285 | 172138 | 262 | High-quality | 96.51 | 0 | (71) |
| Klebsiella phage vB_KpnM_M1‡ | MW448170 | 176329 | 275 | High-quality | 98.85 | 0 | Unpublished |
| SAMN00791912_a1_ct12238‡ | — | 174367 | 272 | High-quality | 97.76 | 0 | (72) |
| uvig_221808‡ | — | 18523 | 17 | Genome-fragment | 10.78 | 0 | (73) |
| uvig_279022‡ | — | 16907 | 33 | Genome-fragment | 9.48 | 0 | (73) |
| uvig_279024‡ | — | 15914 | 37 | Genome-fragment | 8.93 | 0 | (73) |
| uvig_279033‡ | — | 12820 | 25 | Genome-fragment | 7.19 | 0 | (73) |
| uvig_298784‡ | — | 175996 | 274 | High-quality | 98.67 | 0 | (73) |
| uvig_335830‡ | — | 174400 | 272 | High-quality | 97.77 | 0 | (73) |
|  |  |  |  | Quality† | Completeness (%) | Contamination (%) |  |
| uvig_373971‡ | – | 16673 | 24 | Genome-fragment | 7.20 | 22.93 | (73) |
| uvig_510342‡ | – | 18306 | 38 | Genome-fragment | 10.26 | 0 | (73) |
| uvig_510345‡ | – | 15123 | 34 | Genome-fragment | 8.48 | 0 | (73) |
| Zuo_2017_SRR5677802_NODE_1_length_175069_cov_4814.635012‡ | – | 175069 | 272 | High-quality | 100 | 0 | (74) |

\* Annotated using the PHROG hmm database with Prokka.
† Complete, completeness assessed DTR (high-confidence); for all other annotations (i.e. high-quality, genome fragment) completeness was assessed by AAI (high-confidence).
‡ Detected using PhageClouds.

### Analysis of homing endonucleases (HENs) encoded within phage genomes

Genes in all genomes included in the initial ViPTree analysis **(Table 2)** were predicted and annotated using Prokka 1.14.6 (34) using the PHROG (41) dataset. Data were imported into R using Biostrings v2.58.0 (42) and predicted protein names searched for ‘HNH | homing’ to identify HENs encoded with the phage genomes. The HEN sequences were exported in fasta format and imported into Geneious Prime. An MSA was created using Clustal Omega v1.2.3 (options selected: group sequences by similarity, evaluate full distance matrix, 5 refinement iterations). From the MSA, a neighbour-joining tree (Jukes–Cantor) was generated. The tree was uploaded to iToL v6.4.2 (43) for annotation. All data associated with the HEN analysis are available as **Supplementary Material.**

### Comparative genome analyses

PhageClouds uses a gene-network approach to allow rapid searching of ~640,000 phage genomes via a web interface, and includes metagenome-assembled genomes (MAGs) derived from several large-scale metagenome and virome studies (44). Sequences of our initial set of GenBank genomes **(Table 2)** were searched against the PhageClouds database to identify slopekvirus genomes not included in our original analysis, and to determine whether such genomes had been detected in metagenome/virome studies. Results from PhageClouds searches were manually checked to identify a non-redundant set of genomes potentially representing slopekviruses. Quality and completeness of the genomes were determined using CheckV. ViPTree was used to confirm the newly identified genomes fell within the genus *Slopekvirus*.

For all genomes found to be of high quality or complete (*n*=24; **Table 2),** VIRIDIC was used to calculate the intergenomic similarities of the virus sequences (45). The 24 genomes were Prokka-annotated as described above and included in a pangenome analysis (Roary v3.12.0, 95 % identity; (46). Using treeio v1.18.1 (47) and ggtree v3.2.1 (48) a phylogenetic tree was generated from the Roary-generated newick file (accessory_binary_genes.fa.newick), while the binary gene presence/absence file (gene_presence_absence.Rtab) was used to visualize the core and accessory genes identified in the slopekvirus pangenome. For each of the 24 genome sequences, amino acid sequences of core genes with ≥95 % nucleotide identity and ≥70 % coverage (based on comparisons of minimum, average and maximum gene group sizes determined from Roary outputs) were concatenated. These sequences were then used to generate an MSA with MUSCLE v3.8.1551 (49). A maximum likelihood tree was generated from the MSA using PHYML v3.3.20180214 (BLOSUM62, 100 bootstraps; (50), and visualized using iToL v6.4.2. The Prokka-annotated genomes and outputs from Roary are available as **Supplementary Material.**

## RESULTS

### Morphological and genomic characterization of phages isolated on *K. michiganensis*

Two phages (vB_KmiM-2Di and vB_KmiM-4Dii) had been isolated and purified on *K. michiganensis* strains PS_Koxy2 and PS_Koxy4, respectively, during ongoing studies focussed on finding alternatives to antibiotics for treating *Klebsiella*-associated infections (25). In this study, genome sequence data and TEM images were generated for both phages. The genomes were found to be of high quality and free of contamination using CheckV **(Table 2):** the linear genome of vB_KmiM-2Di was 99.34 % complete, comprising 177,200 nt and encoding 275 genes; the linear genome of vB_KmiM-4Dii was 98.02 % complete, comprising 174,857 nt and encoding 271 genes. Neither phage encoded antimicrobial resistance genes. Initial BLAST-based and ViPTree analyses suggested the genomes represented members of the genus *Slopekvirus*. A ViPTree analysis, incorporating all slopekvirus genomes known to us at the time of analysis, confirmed this association **(Fig. 1a).** vB_KmiM-2Di clustered most closely with *Klebsiella virus PMBT1* and *Enterobacter virus Eap3*, whereas vB_KmiM-4Dii was most closely related to *Klebsiella* phage vB_KoM-MeTiny and *Klebsiella* phage vB_KoM-Pickle. ANIm analysis **(Fig. 1b)** showed vB_KmiM-2Di shared ≥99 % ANI with *Klebsiella virus PMBT1*. vB_KmiM-4Dii shared ≥99 % ANI with *Escherichia* phage phT4A, *Klebsiella* phage vB_KoM-MeTiny and *Klebsiella* phage vB_KoM-Pickle.

**Fig. 1.**
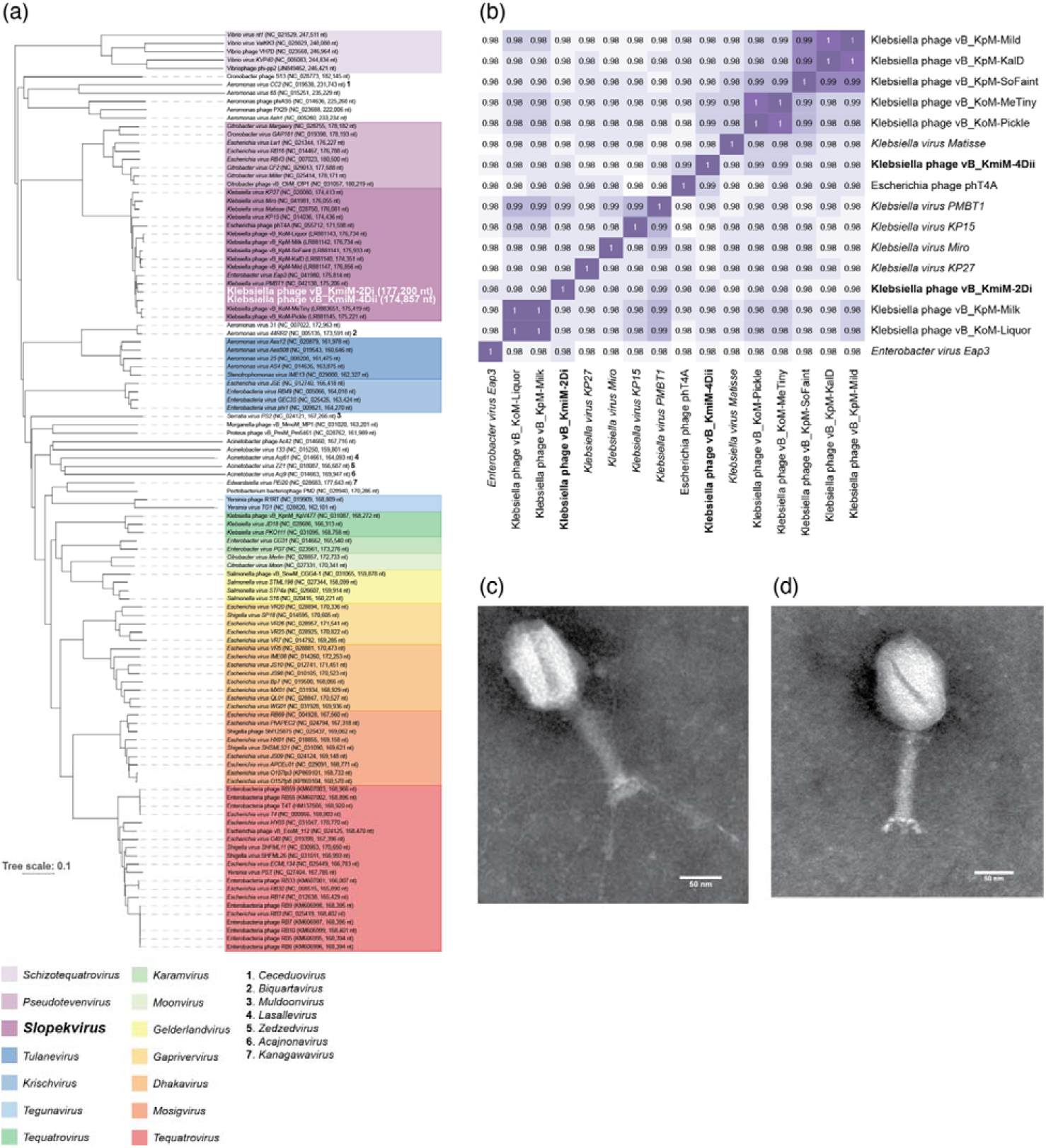
Genomic analyses of slopekviruses *Klebsiella* phage vB_KmiM-2Di and *Klebsiella* phage vB_KmiM-4Dii and related viruses. (a) ViPTree-generated phylogenetic tree of slopekviruses and their closest relatives. (b) ANIm analysis of vB_KmiM-2Di and vB_KmiM-4Dii genomes with members of the genus *Slopekvirus*. (c) Phage vB_KmiM-2Di. (d) Phage vB_KmiM-4Dii. (c, d) Scale bar, 50 nm.

TEM images **(Fig. 1c, d)** showed both phages were myoviruses due to their contractile tails. Phage vB_KmiM-2Di **(Fig. 1c)** had a clearly visible base plate and long tail fibres. The tail and baseplate were recorded at 122.7 nm in length and the capsid diameter was 119.4 nm. The total length of the phage was 242.1 nm. Phage vB_KmiM-4Dii **(Fig. 1d)** had a capsid diameter of 119.7 nm. The length of its tail and baseplate were recorded at 132.1 nm and the total length was 251.8 nm. Three short tail fibres could be seen attached at the bottom of the base plate and short whiskers protruded from the collar under the capsid.

### Identification of clinical and veterinary KoC isolates

*rpoB* gene sequence data were generated for clinical (*n*=59), and veterinary (*n*=49) isolates previously identified as *K. oxytoca* using MALDI-TOF MS and API 20E tests **(Table 1**). These data were compared against the 45 *rpoB* reference allele sequences (released 19 June 2020) available for download from the PubMLST *Klebsiella oxytoca/michiganensis/grimontii* typing database and *rpoB* gene sequences of the type strains of *Klebsiella* (including *Raoultella*) spp. **(Supplementary Fig. 1**). This was done to confirm identity of isolates as we (and others) have previously shown that phenotypic tests and MALDI-TOF are often inadequate for characterization of KoC isolates (10).

Analysis of the sequence data revealed *K. michiganensis* was the most prevalent species represented in both clinical and veterinary isolates (46 % and 43 %, respectively; 27/59 clinical, 21/49 veterinary; **Fig. 2a),** with the *rpoB* gene sequences of all isolates clustering in the clade with the *rpoB* gene sequence of *K. michiganensis* W14^⊤^ **(Supplementary Fig. 1**). *K. oxytoca* was the second most prevalent species represented in the clinical isolates (40 %; 24/59), but only represented 6 % of veterinary isolates (3/49). *K. grimontii* was the second most-common bacterium among the veterinary isolates (24 %; 12/49) and third most-prevalent species in the clinical isolates (12 %; 7/59). *R. ornithinolytica* (12 %; 6/49), *K. huaxiensis* (4 %; 2/49) and *R. terrigena* (2 %; 1/49) were only represented in veterinary isolates. *K. pneumoniae* represented 2 % (1/59) of clinical isolates and 8 % (4/49) of veterinary isolates.

**Fig. 2.**
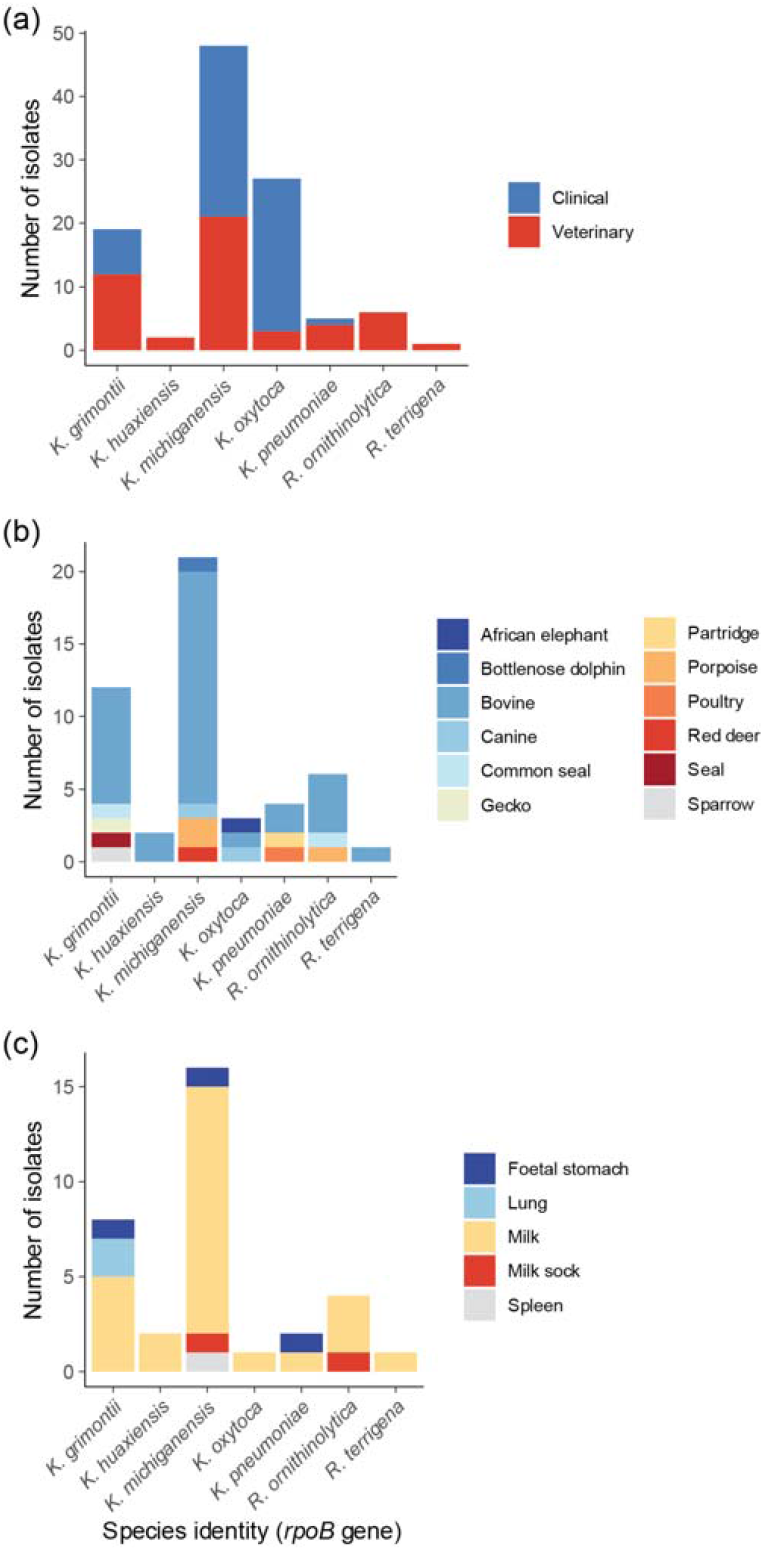
Graphical representations of sources of clinical and veterinary isolates included in this study. (a) Distribution of clinical (*n*=59) and veterinary (*n*=49) isolates among the nine species of bacteria identified in this study. (b) Association of veterinary isolates (*n*=49) recovered from different animals with the nine species of bacteria identified in this study. (c) Association of bovine-associated isolates (*n*=34) with eight different species of bacteria identified in this study.

The majority (73 %; 36/49) of veterinary isolates were of bovine origin, isolated predominantly from milk-related samples (61 %; 30/49) **(Fig. 2b).** While isolates of *K. grimontii* (8/36), *K. huaxiensis (2/36), K. oxytoca* (1/36), *K. pneumoniae* (2/36), *R. ornithinolytica* (4/36) and *R. terrigena* (1/36) had been recovered from milk samples, the majority (44 %; 16/36) of the bovine isolates were *K. michiganensis* **(Fig. 2c).**

### Host range analysis

vB_KmiM-2Di and vB_KmiM-4Dii were screened against clinical (*n*=59) and veterinary (*n*=49) isolates using an agar overlay method **(Table 3).** Phage activity against each strain was recorded as ++, complete lysis; +, hazy lysis; 0, no visible plaques. Depolymerase activity (d) was also recorded.

**Table 3.** Host range analysis of phage vB_KmiM-2di and phage vB_KmiM-4Dii

| Lab ID* | *rpoB* gene-based identity | vB_KmiM-2Di | vB_KmiM-4Dii | Lab ID* | *rpoB* gene-based identity | vB_KmiM-2Di | vB_KmiM-4Dii |
| --- | --- | --- | --- | --- | --- | --- | --- |
| Ko1 | *K. oxytoca* | + | + | GFKo2 | *K. grimontii* | 0 | + |
| Ko2 | *K. oxytoca* | + | + | GFKo3 | *K. grimontii* | 0 | + |
| Ko3 | *K. michiganensis* | + | ++ | GFKo4 | *R. ornithinolytica* | ++ | ++ |
| Ko4 | *K. oxytoca* | + | ++ | GFKo5 | *K. grimontii* | + | + |
| Ko5 | *K. michiganensis* | + | + | GFKo6 | *K. oxytoca* | 0 | 0 |
| Ko6 | *K. oxytoca* | + | + | GFKo7 | *K. michiganensis* | +,d | ++,d |
| Ko7 | *K. oxytoca* | 0 | + | GFKo8 | *K. michiganensis* | 0 | + |
| Ko8 | *K. grimontii* | + | + | GFKo9 | *R. ornithinolytica* | 0 | 0 |
| Ko9 | *K. grimontii* | + | ++ | GFKo10 | *K. michiganensis* | 0 | + |
| Ko10 | *K. michiganensis* | 0 | + | GFKo11 | *K. huaxiensis* | + | + |
| Ko11 | *K. oxytoca* | ++ | ++ | GFKo12 | *K. grimontii* | 0 | 0 |
| Ko12 | *K. oxytoca* | + | ++ | GFKo13 | *K. michiganensis* | 0 | 0 |
| Ko13 | *K. michiganensis* | +,d | +,d | GFKo14 | *K. michiganensis* | + | + |
| Ko14 | *K. michiganensis* | 0 | + | GFKo15 | *K. michiganensis* | + | ++ |
| Ko15 | *K. oxytoca* | + | ++ | GFKo16 | *K. oxytoca* | ++ | ++ |
| Ko16 | *K. grimontii* | ++ | ++ | GFKo17 | *R. ornithinolytica* | 0 | 0 |
| Ko17 | *K. oxytoca* | + | ++ | GFKo18 | *K. michiganensis* | + | + |
| Ko18 | *K. michiganensis* | ++ | ++ | GFKo19 | *K. michiganensis* | + | + |
| Ko19 | *K. oxytoca* | ++ | ++ | GFKo20 | *K. grimontii* | ++ | ++ |
| Ko20 | *K. oxytoca* | + | 0 | GFKo21 | *K. grimontii* | ++ | ++ |
| Ko21 | *K. michiganensis* | + | + | GFKo22 | *K. grimontii* | + | + |
| Ko22 | *K. michiganensis* | + | + | GFKo24 | *R. ornithinolytica* | 0 | + |
| Ko23 | *K. michiganensis* | + | + | GFKo25 | *R. ornithinolytica* | 0 | + |
| Ko24 | *K. michiganensis* | + | + | GFKo26 | *K. grimontii* | + | + |
| Ko25 | *K. oxytoca* | 0 | + | GFKo27 | *K. grimontii* | + | + |
| Ko26 | *K. oxytoca* | ++ | ++ | GFKo28 | *K. michiganensis* | + | ++ |
| Ko27 | *K. grimontii* | + | ++ | GFKo29 | *K. michiganensis* | ++ | + |
| Ko28 | *K. michiganensis* | + | + | GFKo30 | *K. michiganensis* | 0 | 0 |
| Ko29 | *K. michiganensis* | + | + | GFKo31 | *K. michiganensis* | + | + |
| Ko30 | *K. grimontii* | 0 | + | GFKo32 | *K. michiganensis* | 0 | + |
| Ko31 | *K. michiganensis* | + | ++ | GFKo33 | *K. grimontii* | 0 | 0 |
| Ko32 | *K. michiganensis* | + | + | GFKo34 | *K. grimontii* | + | + |
| Ko33 | *K. michiganensis* | 0 | + | GFKo35 | *K. michiganensis* | + | + |
| Ko34 | *K. pneumoniae* | 0 | + | GFKo36 | *K. michiganensis* | + | ++ |
| Ko35 | *K. michiganensis* | + | 0 | GFKo37 | *K. michiganensis* | + | + |
| Ko36 | *K. michiganensis* | 0 | + | GFKo38 | *K. michiganensis* | 0 | ++ |
| Ko37 | *K. oxytoca* | + | + | GFKo39 | *K. grimontii* | ++ | ++ |
| Ko38 | *K. oxytoca* | + | + | GFKo40 | *K. michiganensis* | + | + |
| Ko39 | *K. michiganensis* | + | + | GFKo41 | *K. michiganensis* | 0 | +,d |
| Ko40 | *K. oxytoca* | 0 | 0 | GFKo42 | *K. pneumoniae* | 0 | + |
| Ko41 | *K. michiganensis* | 0 | 0 | GFKo43 | *R. ornithinolytica* | 0 | 0 |
| Ko42 | *K. grimontii* | + | + | GFKo44 | *K. michiganensis* | 0 | 0 |
| Ko43 | *K. michiganensis* | + | + | GFKo45 | *K. michiganensis* | + | + |
| Ko44 | *K. oxytoca* | + | + | GFKo46 | *K. pneumoniae* | 0 | + |
| Ko45 | *K. oxytoca* | + | + | GFKo47 | *K. oxytoca* | 0 | 0 |
| Ko46 | *K. michiganensis* | 0 | 0 | GFKo48 | *K. pneumoniae* | 0 | + |
| Ko47 | *K. oxytoca* | + | + | GFKo49 | *K. pneumoniae* | 0 | 0 |
| Ko48 | *K. oxytoca* | + | + | GFKo50 | *K. huaxiensis* | 0 | 0 |
| Ko49 | *K. michiganensis* | 0 | + | GFKo52 | *R. terrigena* | ++ | ++ |
| Ko50 | *K. michiganensis* | + | + |  |  |  |  |
| Ko51 | *K. grimontii* | + | + |  |  |  |  |
| Ko52 | *K. michiganensis* | + | + |  |  |  |  |
| Ko53 | *K. oxytoca* | + | + |  |  |  |  |
| Ko54 | *K. oxytoca* | 0 | + |  |  |  |  |
| Ko55 | *K. oxytoca* | + | + |  |  |  |  |
| Ko56 | *K. michiganensis* | + | + |  |  |  |  |
| Ko57 | *K. oxytoca* | + | + |  |  |  |  |
| Ko58 | *K. michiganensis* | + | + |  |  |  |  |
| Ko59 | *K. michiganensis* | + | + |  |  |  |  |

\* Ko prefix, clinical isolate; GFKo prefix, veterinary isolate.
† ++, complete lysis; +, hazy lysis; 0, no visible plaques; d, depolymerase activity.

Both phages displayed a broad host range against clinical and veterinary isolates tested. vB_KmiM-2Di showed lytic activity 66 % (71/108) of tested strains (78 %, 46/59 clinical and 51 %, 25/49 veterinary strains, respectively). Whereas vB_KmiM-4Dii showed lytic activity against 84 % (91/108) strains (92 %, 54/59 clinical and 76 %, 37/49 veterinary strains, respectively). The two phages showed lytic activity against one or more strains of *K. michiganensis, K. oxytoca, K. grimontii, R. ornithinolytica, K. huaxiensis* and *R. terrigena*. Additionally, vB_KmiM-4Dii showed activity against *K. pneumoniae*.

The formation of haloes around plaques, indicating depolymerase activity, was observed for both phages against strains Ko13 and GFKo7. vB_KmiM-4Dii also displayed depolymerase activity against strain GFKo41. Neither vB_KmiM-2Di nor vB_KmiM-4Dii was able to infect 17 % (18/108) of the strains tested, the majority (80 %, 12/15) of which were of veterinary origin.

### Analysis of HENs encoded within slopekvirus genomes

It has previously been suggested that differences in host range for members of the genus *Slopekvirus* may be due to HENs encoded within their genomes. These HENs may act as regulators of DNA modification and provide resistance to host restriction enzymes (51). Fifty-four putative HENs were encoded within 13/16 *Slopekvirus* genomes **(Fig. 3).** Several genomes encoded multiple putative HENs, with *Escherichia* phage phT4A (NC_055712) encoding nine. vB_KmiM-2Di encoded five HENs whereas vB_KmiM-4Dii encoded four. Of these, two had possible homologues in both genomes **(Supplementary Fig. 2).** MZ707156_00002 shared homology with MZ707157_00002; 79.7 % pairwise amino acid identity. MZ707156_00164 shared homologywith MZ707157_00163; 100 % pairwise amino acid identity.

**Fig. 3.**
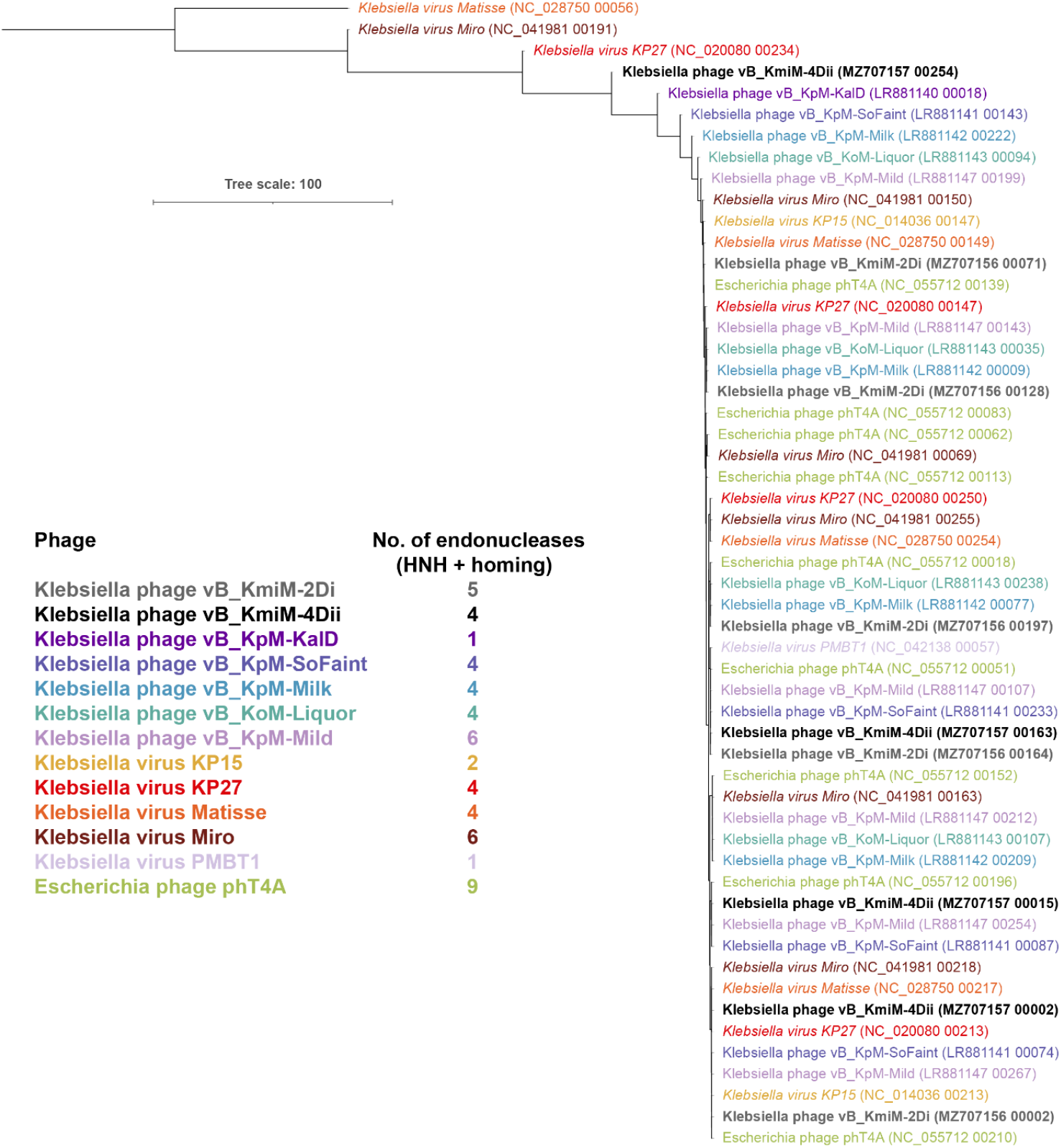
Phylogenetic analysis (neighbour-joining tree) and summation of endonucleases encoded in slopekviruses. In the tree each gene product is coloured based on the phage that encodes it. Scale bar, average number of amino acid substitutions per position. The alignment, distance matrix and newick file associated with the analysis are available from figshare as **Supplementary Material.**

### Diversity of slopekviruses in GenBank and metagenomic data

PhageClouds was used to identify slopekviruses not included in our initial scoping of viruses related to phages vB_KmiM-2Di and vB_KmiM-4Dii. Each PhageClouds search generated 33 hits. **Supplementary Table 1** provides example output for a search done using the genome sequence of phage vB_KmiM-2Di. Seventeen of the 33 hits represented sequences included in our initial phylogenetic analysis **(Fig. 1a),** six represented isolated phages not included in our initial list of viruses and 11 represented MAGs recovered from virome studies **(Table 2).** ViPTree analysis showed all 17 viruses were related to the genus *Slopekvirus* (data not shown). CheckV analysis showed four of the GenBank and four of the MAG sequences represented complete or high-quality phage genomes **(Table 2).**

Our initial analyses **(Fig. 1, Supplementary Fig. 3)** showed that slopekvirus genomes shared high sequence similarity, with the HENs contributing to their diversity. Consequently, we used all 24 high-quality/complete slopekvirus genomes identified in this study **(Table 2)** in a pangenome analysis to determine whether core genes could be identified within the genus *Slopekvirus* **(Fig. 4).** It has been recommended that sequence and similarity coverage of proteins are set to >30 % identity and 50 % coverage, respectively, for genus-level phage-based pangenome studies (52). However, for the genus *Slopekvirus* we found these criteria were too lax (not shown). Our Roary-based pangenome analysis run at 95 % identity identified 155 core genes in a total pangenome of 425 genes **(Fig. 4a, b; Supplementary Table 2).** The pangenome was open, as the plot showing the total number of genes is not asymptotic **(Fig. 4c).** Filtering the core genes based on comparisons of minimum, maximum and average group nucleotide coverage **(Supplementary Table 2)** identified 148 core genes had ≥95 % identity and ≥70 % coverage. Given that the 24 genomes encoded a mean of 272 genes (+/- 5 genes) each, these core genes represented 54 % (148/272) of the total genome content of the slopekviruses. The majority (61/148, 41 %) of these genes were predicted to encode hypothetical proteins, with baseplate wedge subunit proteins (6/148, 4 %), proteins of unknown function (4/148, 3 %), tail tube (3/148, 2%) and RIIB lysis inhibitor, major head, head scaffolding, clamp loader of DNA polymerase, baseplate hub and 5’-3’ deoxyribonucleotidase proteins (2/148, 1 % each) making the greatest contribution to the core genes **(Supplementary Fig. 4).** As expected, the HENs contributed to the accessory genes.

**Fig. 4.**
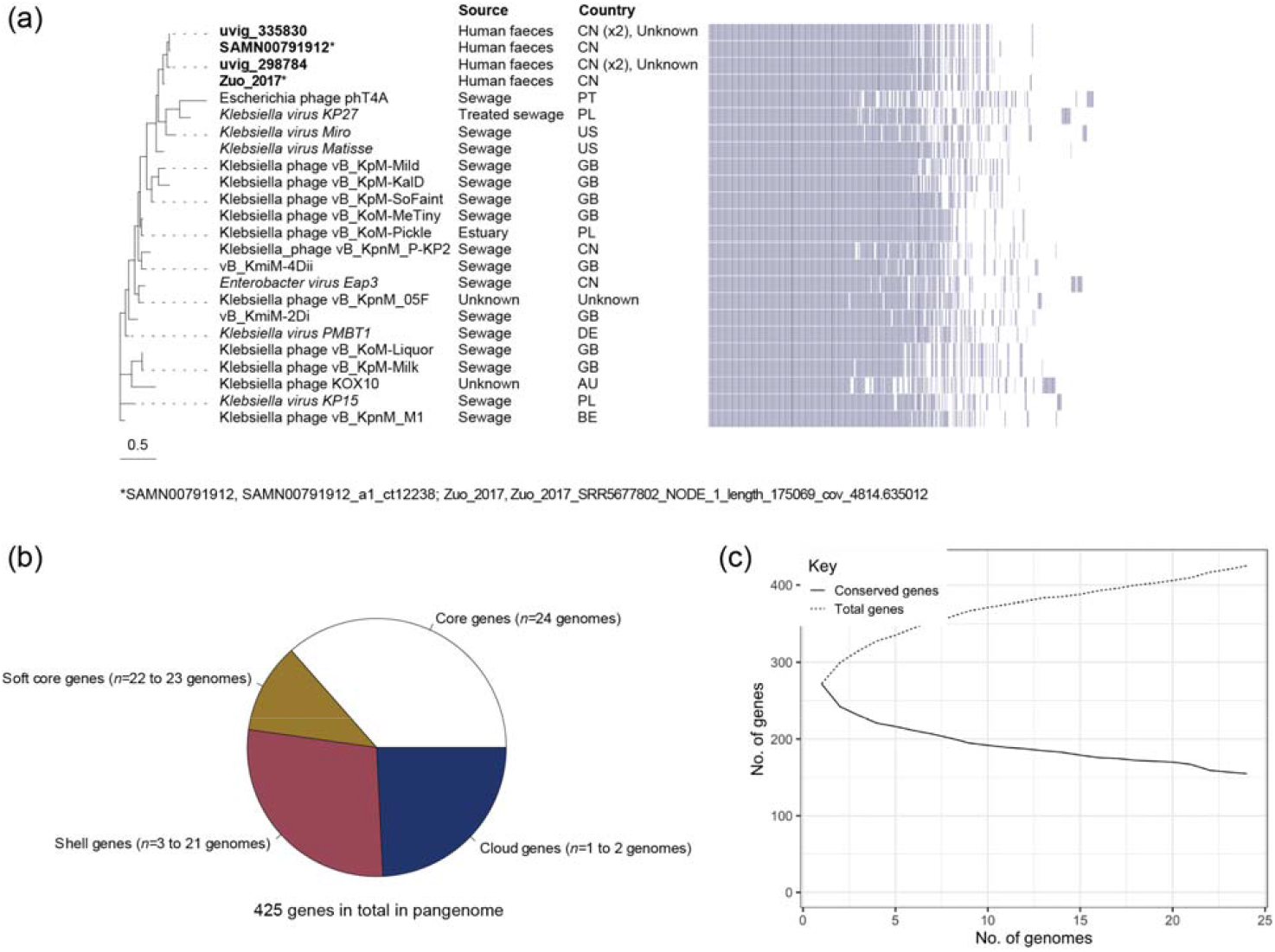
Visualization of pangenome data for 24 slopekvirus genomes. (a) Roary matrix of the 425 genes representing the total pangenome. AU, Australia; BE, Belgium; CN, China; GB, Great Britain; DE, Denmark; PL, Poland; PT, Portugal; US, United State of America. (b) Pie chart showing the contribution of the 425 genes to the pangenome. (c) Plots showing how the number of total genes (dotted line) and conserved genes (dashed line) changes as more genomes are added to the slopekvirus pangenome.

If two phage genome sequences, tested reciprocally, are more than 95 % identical at the nucleotide level over their full genome length, they are assigned to the same species (52). We, therefore, sought to determine the species diversity within the genus *Slopekvirus*. VIRIDIC analysis of the 24 genome sequences included in our pangenome analysis suggested there were eight species represented within the genus **(Supplementary Table 3),** though bidirectional hierarchical clustering of these data showed no obvious clustering of the species **(Fig. 5a).** An MSA of the concatenated protein sequences for the 148 core genes (available as **Supplementary Material)** showed the total alignment of 32,904 aa shared a minimum identity of 97.16 % across all genomes **(Supplementary Table 4).** Maximum-likelihood phylogenetic analysis **(Fig. 5b)** of these concatenated sequences did not support species separation, nor did clustering of the accessory gene (binary) data in the pangenome analysis **(Fig. 4a).**

**Fig. 5.**
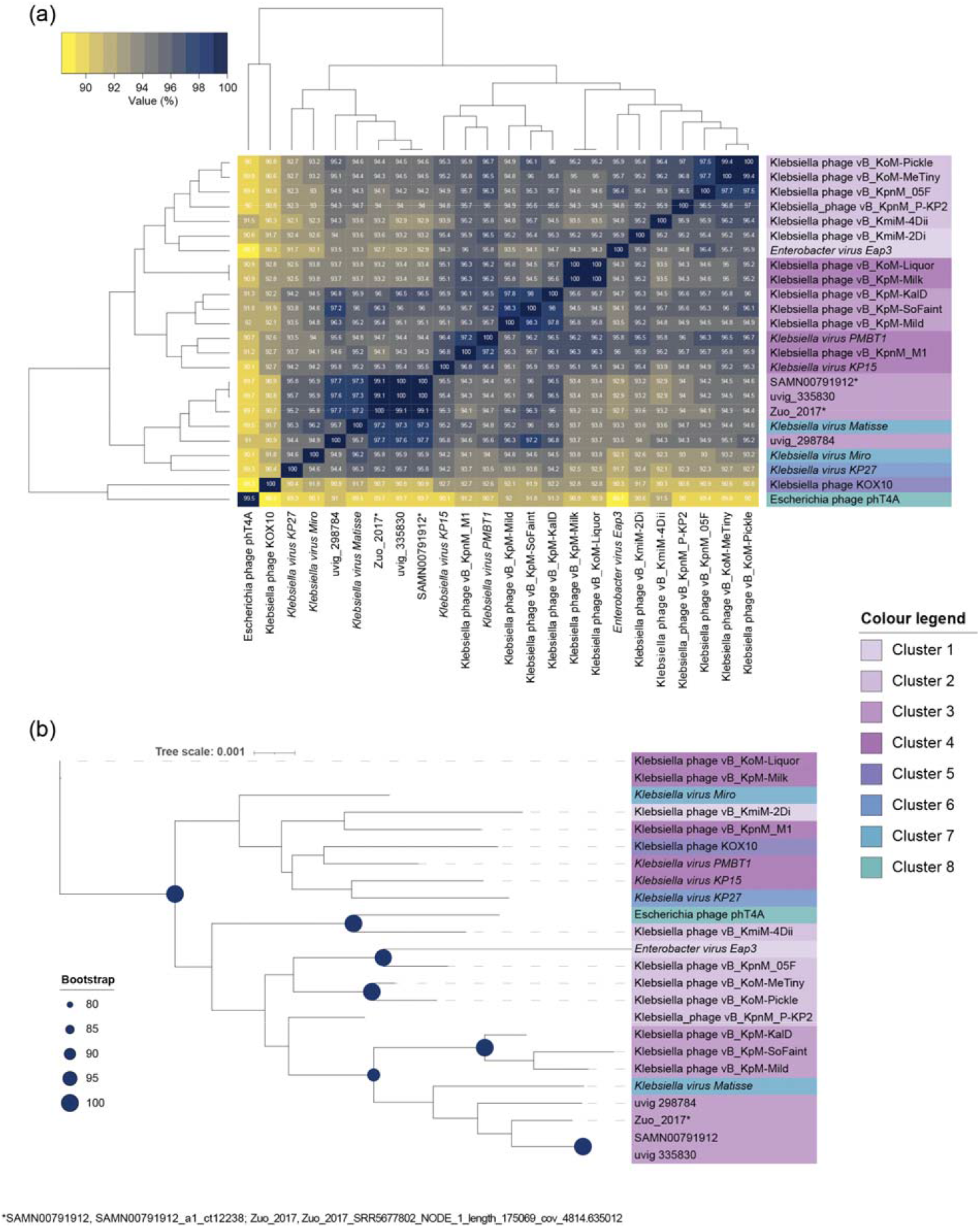
Determination of whether distinct species are represented with the genus *Slopekvirus*. (a) Bidirectional clustering heatmap visualizing VIRIDIC-generated similarity matrix for the 24 slopekvirus genomes. (b) Phylogenetic tree (maximum likelihood) generated by concatenating the amino acid sequences of the 148 core genes from the pangenome that had ≥95 % identity and ≥70 % coverage across the 24 genomes. Bootstrap values are presented as a percentage of 100 replications. (a, b) Colouring of the data corresponds to the eight species clusters predicted by VIRIDIC.

## DISCUSSION

In this study we characterized the morphology and genomes of two phages (vB_KmiM-2Di and vB_KmiM-4Dii) we had isolated on two strains (PS_Koxy2 and PS_Koxy4, respectively) of multidrug-resistant *K. michiganensis* (10,25). Both phages were found to belong to the genus *Slopekvirus* and had the myoviridae-type morphology consistent with other known slopekviruses **(Fig. 1).** The host ranges of these phages were determined on a collection of clinical (*n*=59) and veterinary (*n*=49) bacteria that had been originally identified as belonging to the KoC **(Table 1**). Both phages were found to have broad host ranges **(Table 3).** This feature of the slopekviruses is thought to be due to the number of HENs encoded within their genomes **(Fig. 3).** Searches among ~640,000 phage genomes, using PhageClouds, allowed us to identify slopekviruses within metagenome and virome datasets. We also characterized the genomes of our phages and those of their closest relatives, undertaking a pangenome analysis to better understand the genomic diversity of slopekviruses.

### Identification of members of the KoC from clinical and veterinary sources

Despite our clinical and veterinary isolates being presumptively identified as *K. oxytoca* by MALDI-TOF MS and API 20E profiling, *K. oxytoca* only represented 25 % (27/108) of the isolates identified by *rpoB* gene sequence analysis. While we have previously highlighted that phenotypic assays and MALDI-TOF MS frequently do not allow accurate identification of clinical isolates of the KoC (10,53), this is the first time we have encountered issues identifying veterinary KoC isolates. Veterinary MALDI-TOF MS databases should be updated to include relevant reference spectra to allow differentiation of KoC species (1)

The *rpoB* gene-based sequence analysis identified *K. michiganensis* as the most common isolate from both clinical and veterinary samples (27/59 and 21/49 respectively). This supports our previous findings with respect to *K. michiganensis* being more clinically relevant than *K. oxytoca* (53). Gómez *et al.* also used *rpoB* gene sequence analysis to identify commensal, community-acquired and neonatal intensive care unit *Klebsiella* spp. isolates. Routine biochemical testing identified 21 *K. oxytoca;* however, *rpoB* gene analysis identified them as *K. michiganensis* (16/21), *K. grimontii* (5/21) and *K. pneumoniae* (1/21) (54). Taken together with our work, these findings reinforce the clinical relevance of *K. michiganensis* and its historic underappreciation as a human pathogen due to misidentification.

In recent years, *K. michiganensis* has been isolated from a diverse range of animals: farm animals (cows, poultry and pigs) (13,55), companion animals (cat, dog and horses) (12,55) and other animals including hedgehogs, guineapigs, mice, fruit bats, turtles and invertebrates (13,55–58). This is from using tools capable of discriminating between members of the KoC, but further work is needed to assess the true clinical and epidemiological significance of *K. michiganensis* in animals. Furthermore, many of these studies are retrospective, using accurate but time- and resource-intensive methods that would not be practical for rapid clinical or veterinary diagnosis. It is notable that 16/21 of our *K. michiganensis* veterinary isolates had been recovered from bovine milk samples **(Table 1**; **Fig. 2c).** Whether these isolates encode genetic determinants specific to mastitis, as observed for an acquired *lac* operon in bovine-associated *K. pneumoniae* (59), will be determined when we analyse genome sequence data for them in the future, with results to be reported elsewhere.

*K. grimontii* has previously been isolated from animals in Germany (cattle/milk, rabbit, pig, sheep, dog, pig, tortoise, hedgehog, roe deer) (13), and during a longitudinal study undertaken in Pavia, Italy (sheep, horse, fly, cattle, pig, cat, duck, turtle, dog, cockroach, wasp, chicken) (55). We found *K. grimontii* in seal (*n*=2), bovine (*n*=8), gecko (*n*=1) and sparrow (*n*=1) samples collected in the UK. *K. grimontii* appears to be the second most-common KoC species of veterinary relevance.

*K. huaxiensis* was originally described based on one isolate recovered from human urine in China (16). Since then, the bacterium has been isolated from cow and human faeces in Italy (1), and from cows (*n*=4), water (*n*=2), a horse (*n*=1) and hospital carriage (*n*=1) (55). In this study we identified two isolates (GFKo11, GFKo50) from bovine milk collected in Scotland. As for other members of the KoC, more work is needed to determine the wider relevance of *K. huaxiensis* to veterinary infections.

### Host range determination for phages vB_KmiM-2Di and vB_KmiM-4Dii

We found that phages vB_KmiM-2Di and vB_KmiM-4Dii both exhibited broad host ranges against several different species of bacteria **(Table 3).** This includes isolates of *K. pneumoniae, K. michiganensis, K. oxytoca, K. grimontii, R. ornithinolytica, K. huaxiensis* and *R. terrigena*. Ordinarily, phage host range is narrow, sometimes down to the strain level. However broad-host-range phages are reported in the literature (60–62) and phages with extended host ranges have been identified within the genus *Slopekvirus* (21,38). For example, phage vB_KoM-MeTiny, which is genetically similar to vB_KmiM-2Di and vB_KmiM-4Dii (95.2 % and 96.2 % respectively; VIRIDIC), is reported to form plaques on *K. michiganensis, K. oxytoca, K. pneumoniae, K. variicola* and *K. quasipneumoniae*. These findings suggest bacteriophage belonging to the genus *Slopekvirus* are useful as potential therapeutics owing to their broad host ranges and lack of antimicrobial resistance genes (21,38), in agreement with others who have worked with slopekviruses (21,32).

Despite a high level of sequence identity between vB_KmiM-2Di and vB_KmiM-4Dii (95.25 %; VIRIDIC) differences in host range were observed between the two phages. Maciejewska *et al.* previously characterised two slopekviruses (vB_KpnM_KP15 and vB_KpnM_KP27) that exhibit broad lysis against *Klebsiella* spp. (63). They also noted differences in the host ranges of the two phages, despite a high-level of DNA identity (94.2 %; our VIRIDIC analysis). The authors suggested the discord in host range may be due to the presence of two HENs encoded within the genome of vB_KpnM_KP27, both of which are absent from vB_KpnM_KP15. HENs are site-specific DNA endonucleases that function as mobile genetic elements by catalysing a double-strand break at specific DNA target sites in a recipient genome that lacks the endonuclease. The double-strand break is then repaired by homologous recombination using the allele containing the HEN as the template. The result is the incorporation of the HEN into the cleavage site. Maciejewska *et al.* hypothesised that HENs encoded within vB_KpnM_KP27 (YP_007348875.1 and YP_007348891.1) may act as regulators of DNA modification by splicing events near DNA modification genes located in their close vicinity. Such events may result in protection against host restriction enzymes and subsequently modulate host range. We identified a homologue of YP_007348875.1 (86 % pairwise amino acid identity with MZ707157_00254) and its associated flanking region in the genome of vB_KmiM-4Dii that is absent from vB_KmiM-2Di **(Supplementary Fig. 5).** The second HEN identified in KP27 (YP_007348891.1) was absent from both vB_KmiM-2Di and vB_KmiM-4Dii. The presence of YP_007348875.1 in the genome of vB_KmiM-4Dii may help explain its slightly broader host range compared to vB_KmiM-2Di.

We also conducted a search for other HENs encoded in other slopekvirus genomes. Despite a high degree of genetic conservation across the genus **(Fig. 4, Fig. 5),** there was a high level of divergence with respect to the number and type of HENs encoded across the 16 genomes and between vB_KmiM-2Di and vB_KmiM-4Dii. We found that vB_KmiM-2Di encodes five putative HENs, whereas vB_KmiM-4Dii encodes four. Of note was a HEN identified in the genomes of both vB_KmiM-2Di and vB_KmiM-4Dii (MZ707156_00164 and MZ707157_00163, respectively). This HEN was found immediately upstream of a CDS encoding a putative SbcC-like subunit of a predicted palindrome-specific endonuclease **(Supplementary Fig. 6).** This may again be a case of a HEN splicing event near a gene involved in DNA modification, with the result influencing host range. These findings lend support to the theory that HENs may influence host range and account for the differences in host lysis observed between vB_KmiM-2Di and vB_KmiM-4Dii.

It has been noted by others (21,64) that phages with highly similar sequences may have altered lytic spectra due to selection pressures applied by different hosts used for propagation. In the current study, *K. michiganensis* strains PS_Koxy2 and PS_Koxy4 were used for propagation of vB_KmiM-2Di and vB_KmiM-4Dii, respectively. These two strains have been extensively characterised both genomically and phenotypically and have been shown to share >99.9 % ANI and the same multi-locus sequence type (10). Therefore, we conclude that any differences observed with respect to host range are unlikely to be a consequence of the host used for propagation.

### Genetic diversity of slopekviruses: implications for phage taxonomy

Our Roary-based pangenome analysis of 24 high-quality/complete slopekvirus genomes showed the genus *Slopekvirus* comprises phages with highly conserved genomes, with the core genome (determined using ≥95 % identity and ≥70 % coverage) representing over half of the total genome content of the genus **(Fig. 4).** The recommended cut-off criteria (>30 % identity, 50 % coverage (52)) for genus-level phage pangenome analysis was found to be inappropriate for use with the genus *Slopekvirus*. Results from our analysis (along with other unpublished work from our laboratory) suggest wider-ranging studies are required to determine appropriate recommended identity and coverage cut-off values for use in phage-based pangenome studies at any taxonomic level.

The use of VIRIDIC (and other ANI tools) to assign phage that share ≥95 % reciprocal sequence identity at the nucleotide level over their full genome length to the same species is also questioned (52). Our VIRIDIC analysis of 24 high-quality/complete slopekvirus genomes suggested our dataset represented eight different species **(Fig. 5).** However, phylogenetic analysis based on the protein sequences of 148 core genes (encoding 32,904 aa) did not support separation of the genus into eight species. ANI analyses must be supplemented with phylogenetic analyses to provide robust evidence to support ‘species’ designations within phage genera, as recommended for bacterial and archaeal taxonomy (65).

## AUTHOR STATEMENTS

### Authors and contributors

Conceptualisation: PS, LH, DN. Data curation: all authors. Formal analysis: TSZ, PS, ALM, GF, LH, DN. Funding acquisition: PS, ALM, LH. Investigation: TSZ, PS, ALM, LH, DN. Methodology: ALM, LH, DN. Resources: GF, ALM, LH, DN. Supervision: ALM, LH, DN. Visualisation: TSZ, LH, DN. Writing – original draft: TSZ, LH, DN. Writing – reviewing and editing: all authors.

### Ethical approval

The study of anonymised clinical isolates provided by the Nottingham University Hospitals NHS Trust (NUH) Pathogen Bank was approved by NUH Research and Innovation (19MI001).

## Acknowledgements

This work used computing resources funded by the Research Contingency Fund of the Department of Biosciences, Nottingham Trent University (NTU). TSZ completed this work as part of an MRes degree at NTU. We thank Andrew Millard (University of Leicester) for providing us with the Prokka-formatted PHROG dataset for annotation of phage genomes.

## Abbreviations

AAHC: antibiotic-associated haemorrhagic colitis
AMR: antimicrobial resistance
ANI: average nucleotide identity
HEN: homing endonuclease
KoC: *Klebsiella oxytoca* complex
MAG: metagenome-assembled genome
MALDI-TOF MS: matrix-assisted laser desorption/ionisation-time of flight mass spectrometry
MLST: multi-locus sequence typing
MSA: multiple-sequence alignment
NA: nutrient agar
NB: nutrient broth
TEM: transmission electron microscopy
UTI: urinary tract infection

---

**Supplementary Fig. 1.**
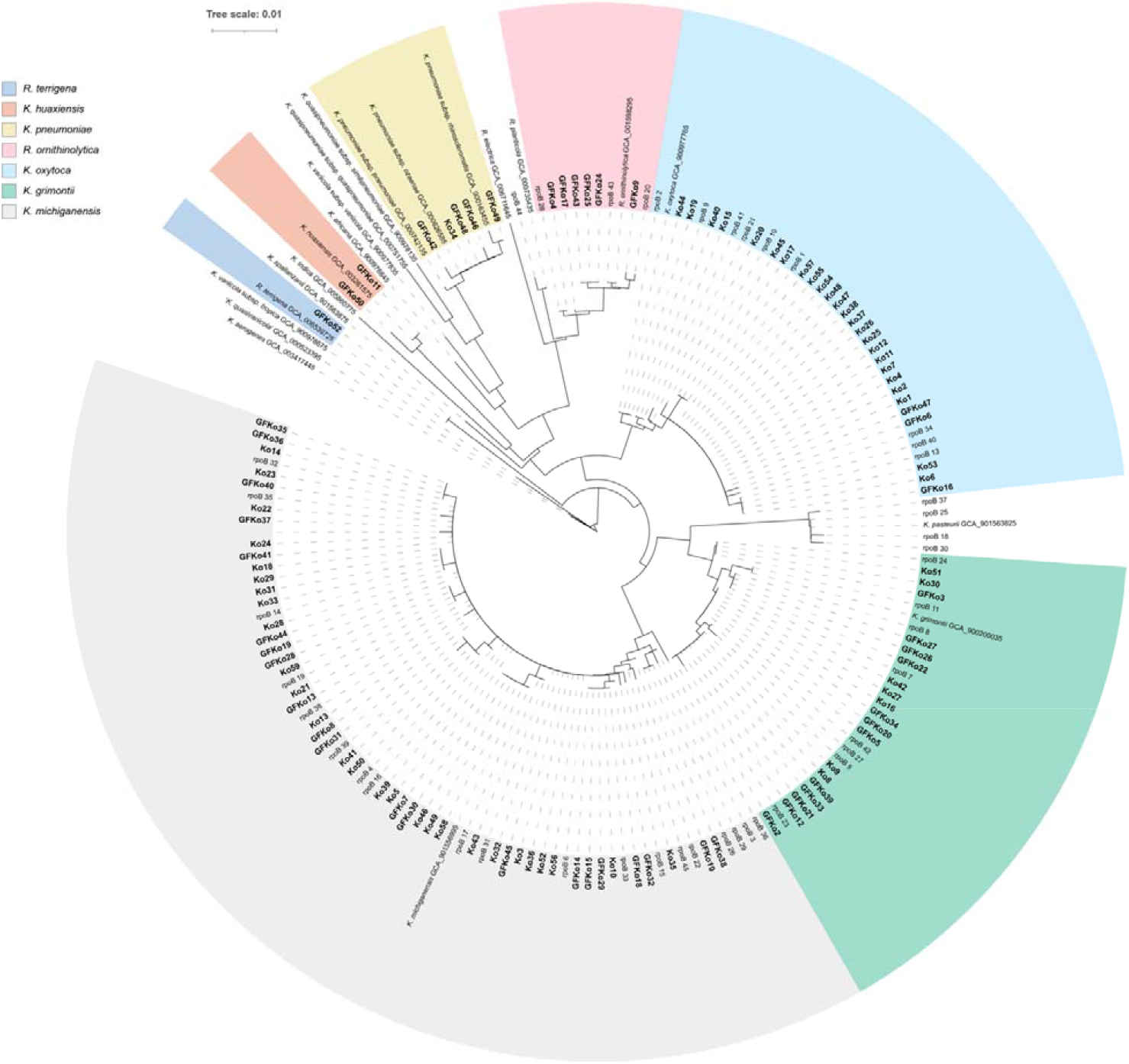
Phylogenetic tree (neighbour joining) showing relationship of clinical (*n*=59) and veterinary (*n*=49) isolates with other *Klebsiella* (including *Raoultella*) spp. Accession numbers shown for type species of *Klebsiella* spp. are for genome sequences from which *rpoB* gene sequences were extracted. The tree was rooted using the *rpoB* gene sequence of *K. aerogenes* as an outgroup. Scale bar, average number of nucleotide substitutions per position.

**Supplementary Fig. 2.**
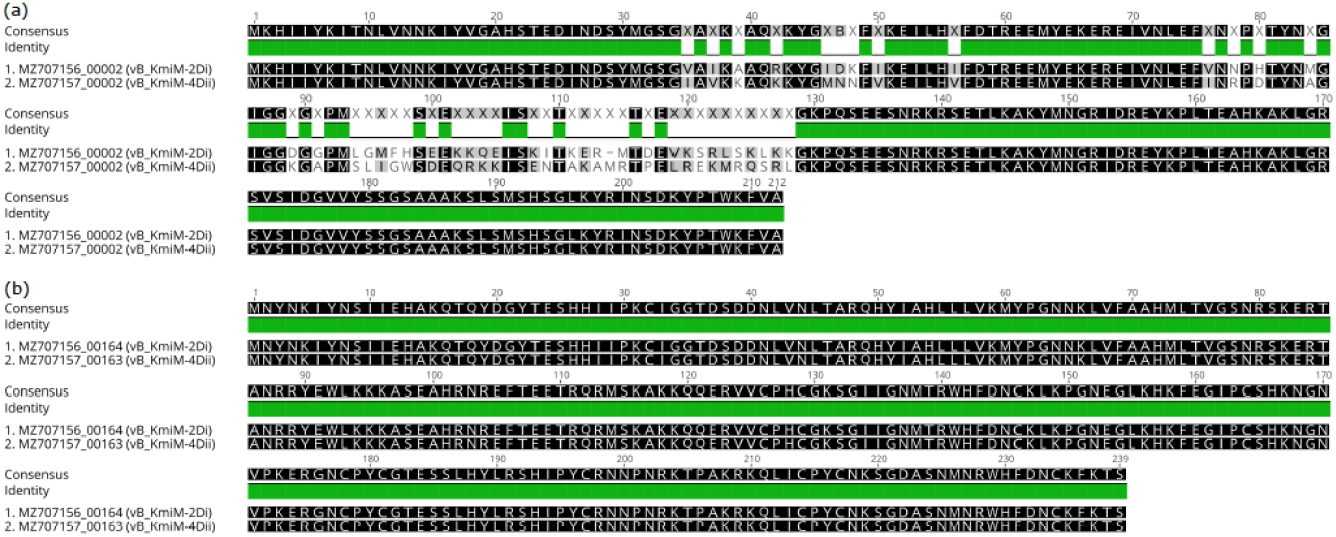
Amino acid alignments of homologous HENs identified in vB_KmiM-2Di and vB_KmiM-4Dii. (a) MZ707156_00002 homologous with MZ707157_00002; 79.7 % pairwise amino acid identity. (b) MZ707156_00164 homologous with MZ707157_00163; 100 % pairwise amino acid identity.

**Supplementary Fig. 3.**
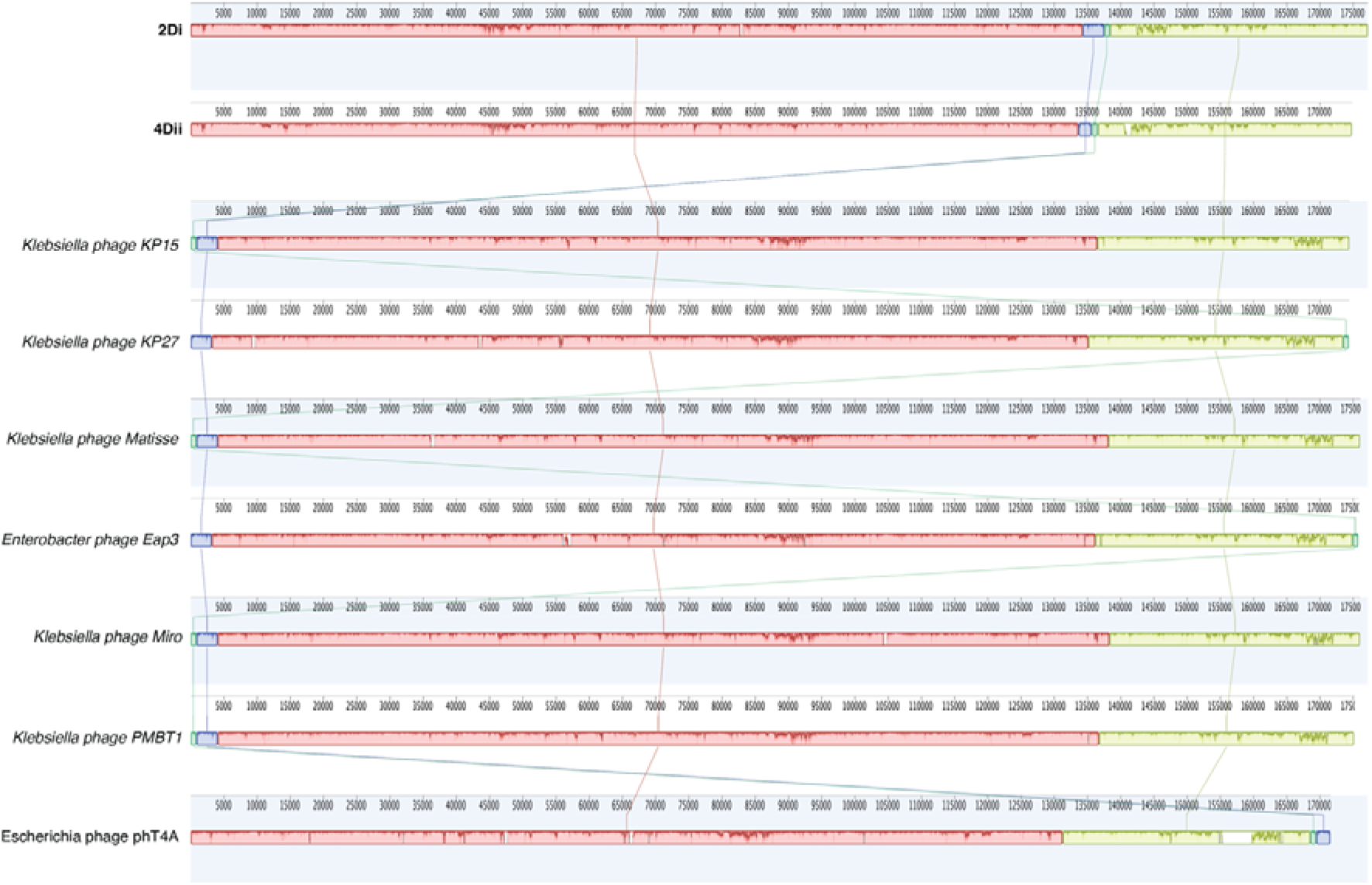
Mauve alignment of phages vB_KmiM-2Di and vB_KmiM-4Dii with representatives of the genus *Slopekvirus*. The alignment shows that the genomes of the phages are highly similar. Visual inspection revealed the HENs contributed to the diversity of the genomes.

**Supplementary Fig. 4.**
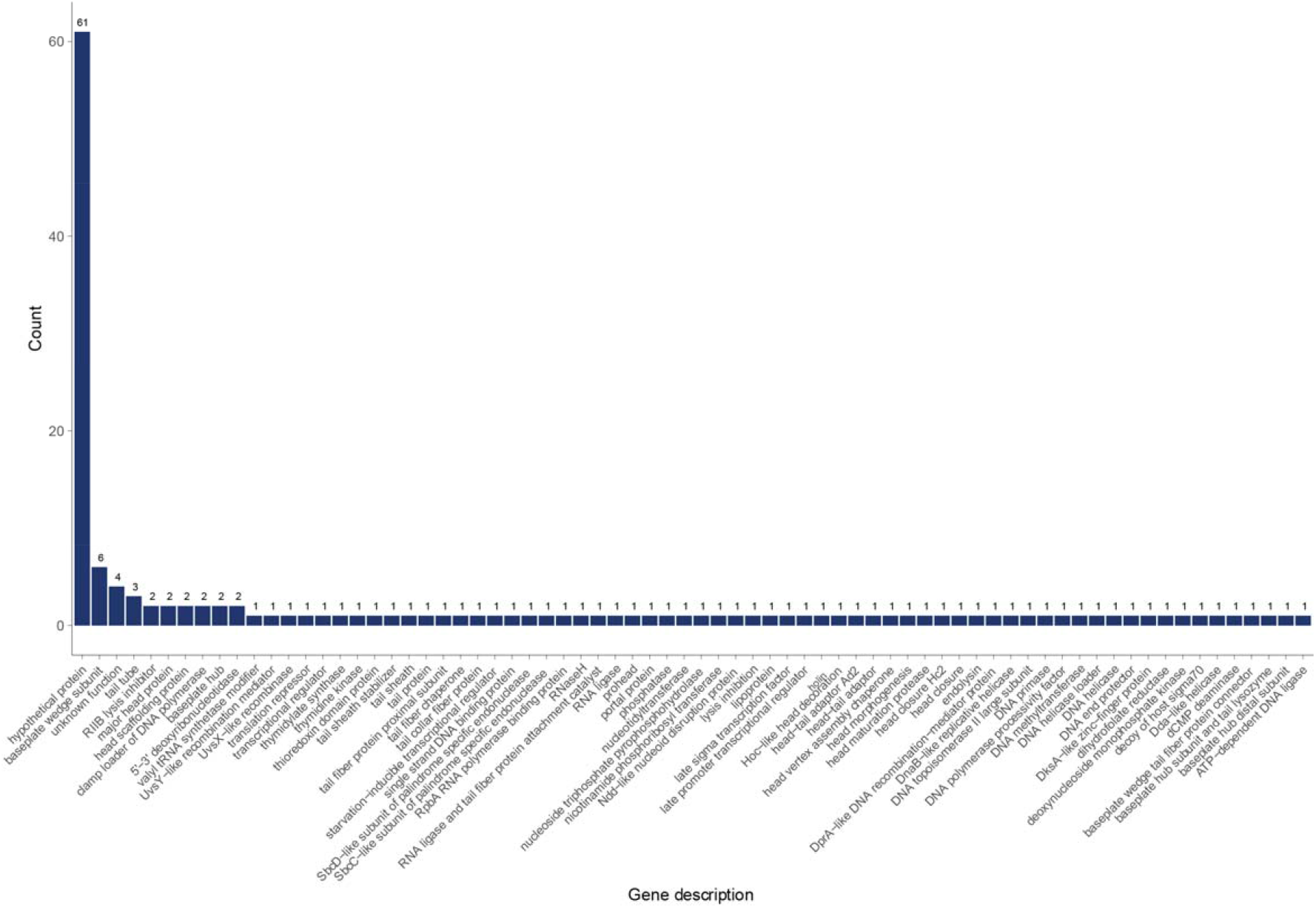
Summary of functions predicted for 148 core genes identified in the slopekvirus pangenome analysis.

**Supplementary Fig. 5.**
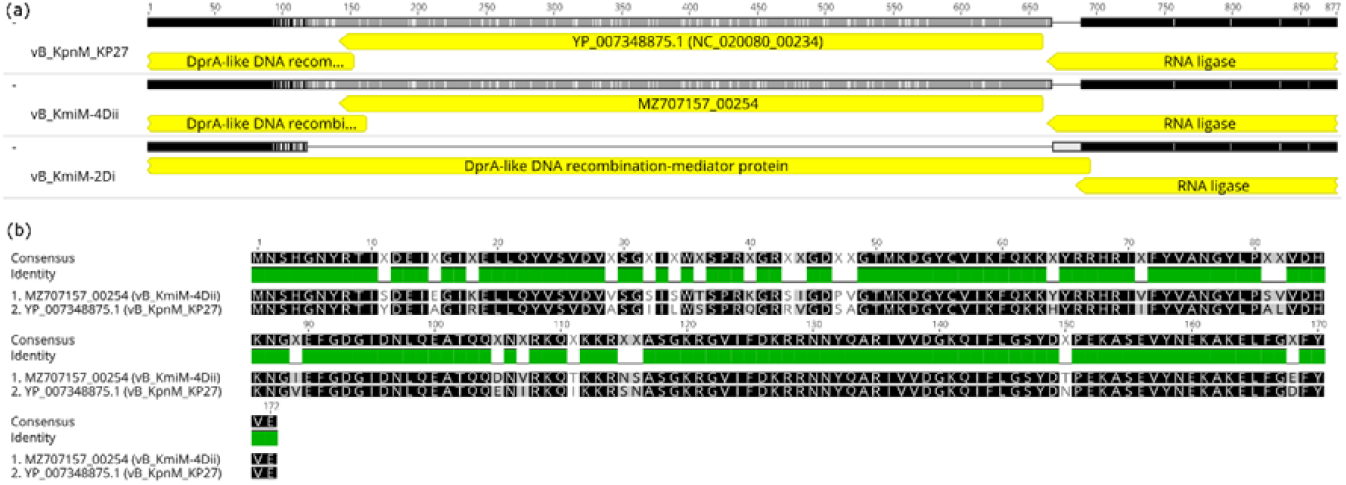
Analysis of HEN YP_007348875.1 originally identified in vB_KpnM_KP27. (a) Genome alignment of vB_KpnM_KP27 with vB_KmiM-2Di and vB_KmiM-4Dii revealed the presence of a potentially homologous gene in vB_KmiM-4Dii that was absent in vB_KmiM-2Di. (b) Amino acid alignment of two translated sequences showed 86 % pairwise amino acid identity between YP_007348875.1 and MZ707157_00254.

**Supplementary Fig. 6.**
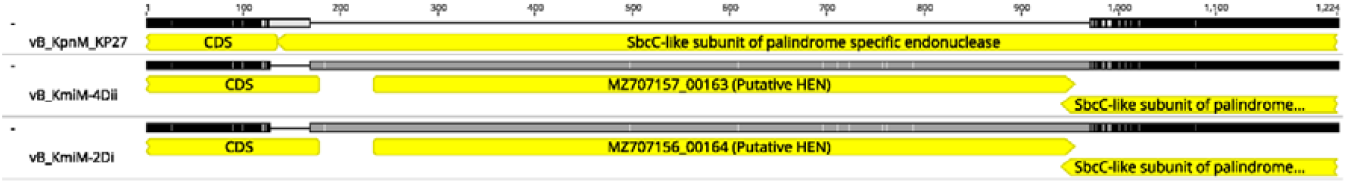
Genomic regions encoding putative homologous HENs in vB_KmiM-2Di and vB_KmiM-4Dii. These HENs were found immediately upstream of a CDS encoding a putative SbcC-like subunit of a predicted palindrome-specific endonuclease. No putative homologue was identified in the genome of vB_KpnM_KP27.

